# Distortion rules: audibility creation in the mosquito ear

**DOI:** 10.1101/2025.09.05.674403

**Authors:** Alexandros C Alampounti, Marcos Georgiades, Justin Faber, Judit Bagi, Marta Andrés, Dolores Bozovic, Joerg T Albert

## Abstract

Hearing begins when ears convert sound into neuronal excitation. The hearing organs of male *Anopheles* mosquitoes rank amongst the most sensitive ears evolved. Males use them to detect faint female flight tones within noisy mating swarms, but female flight tones largely lie above the male’s hearing range. We report a mechanism by which inaudible higher-frequency primary tones become audible through intrinsically generated, lower-frequency distortion tones. Distortion-evoked responses show ∼10-times greater sensitivity and 45% higher temporal precision than those of same-frequency primary tones. Moreover, self-sustained oscillations of the male’s ear dramatically reshape auditory frequency tuning and neuronal recruitment. By one mechanism, mosquito ears thus create audible signals and isolate these from the most salient frequencies of background noise. In audiology, distortions are used for diagnostics; their biological relevance, however, is unknown. Our results suggest that distortions are not mere by-products of the auditory process, but integral to its signaling logic.

## Main Text

All neural systems could be described as networks of interconnected sender-receiver chains, which continually produce and respond to specific signals. Specificity arises when the timing and frequency structure of the incoming signals aligns with the tuning properties of their corresponding receivers (*1*). Yet, the properties of most receivers are only partly known, and signals change as they travel along the sender-receiver chains. Multiple essential nonlinearities (*2*) linked to signal transduction, transmission or transformation distort the signals’ waveforms. The resulting distortion products (DPs) represent newly introduced frequency components — predictable by mathematical rules and tightly bound to the original input stimuli. In hearing, they are well known and widely exploited for clinical diagnostics(*3*), but their roles in sensory perception and neural encoding are unknown.

The malaria mosquito *Anopheles gambiae* offers a powerful model to probe these questions. Males rely on hearing to locate females mid-flight (*4*), a task demanding high sensitivity amid the noise of swarming conspecifics. Although the male’s flagellar sound receiver is mechanically well tuned to female wingbeat frequencies, these fall outside the tuning range of his auditory nerve, revealing a mismatch between mechanical and neural sensitivity. It has been proposed that nonlinear interactions between flight tones generate distortion products that fall within the neural hearing range and that these serve as behavioral cues (*5–10*). In addition to this two-tone distortion, the flagellar ears of *Anopheles* males can generate large-amplitude, near-sinusoidal self-sustained oscillations (SSOs) (*11*, *12*), thereby potentially adding a third component to the tonal mix.

Using laser vibrometry and nerve recordings, we simultaneously captured the system’s mechanical input and neural output. We investigated how distortions emerge, how they reshape neural responses, and how they contribute to audibility.

## Results

### Tonal dissection of an auditory black box

To analyse the response behaviour of the male *Anopheles* ear we tested tonal combinations for distinct, ecologically relevant states (Fig. 1A,B). One-tone-stimuli reflected a flying mosquito - male or female (simulated fly-by tones *f*_1_ = 100 − 1000 Hz) - passing by a stationary, *non-flying* male. Two-tone-stimuli reflected the same mosquito passing by a now *flying* male (*f*_2_=540 Hz).

**Fig. 1.**
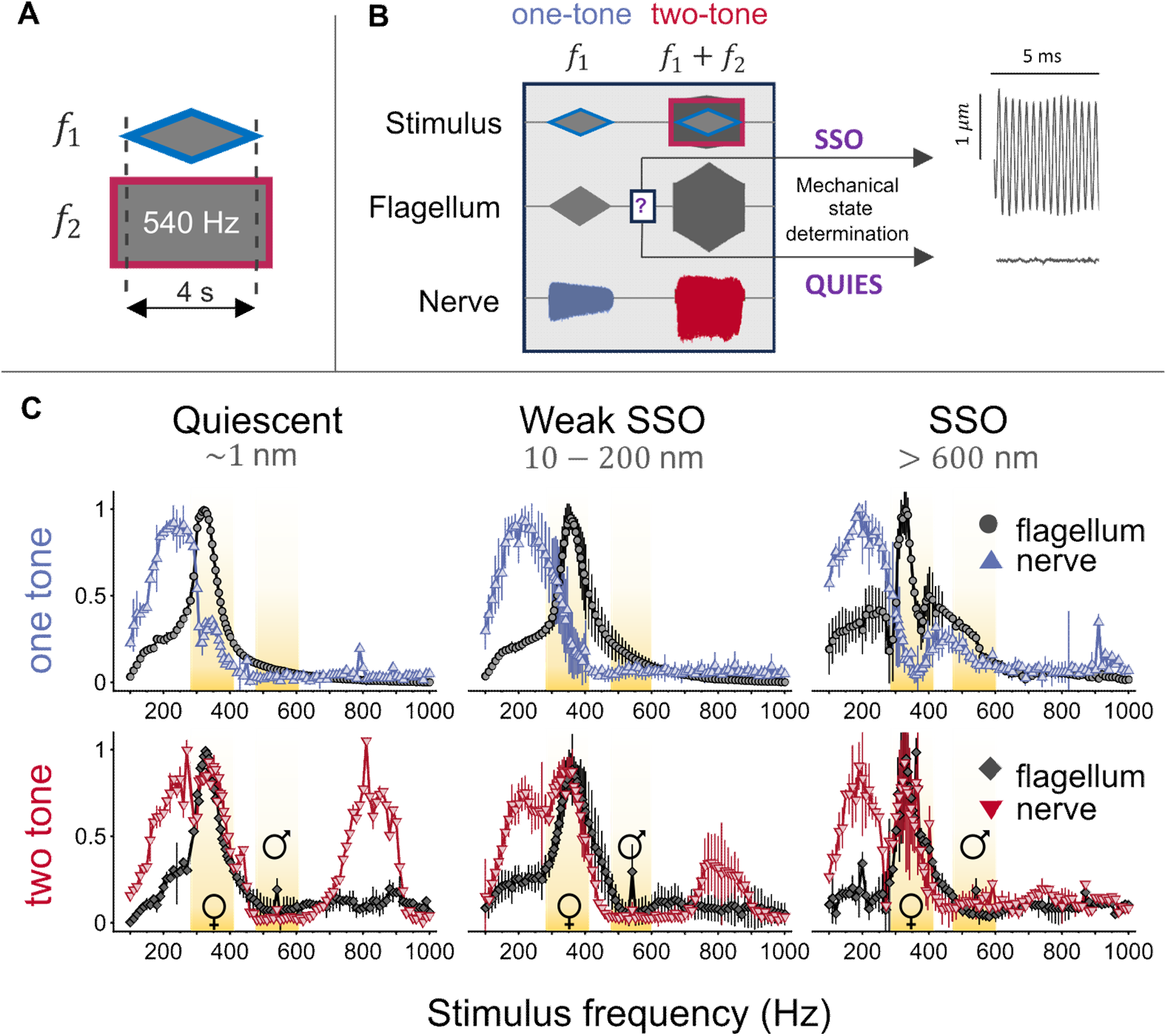
Frequency response functions of male flagellar receiver and auditory nerve for one- and two-tone stimulations across different oscillatory states. **(A)** Two stimulus frequencies, *f*_1_ and *f*_2_, were used either in isolation (‘one-tone’) or combined (‘two-tone’); *f*_1_ was varied whereas *f*_2_was fixed at 540 Hz. **(B)** Experimental design: stimuli, flagellar displacements and compound action potential responses from the antennal nerve were recorded, one-tone and two-tone stimuli were played in sequence separated by a 2s gap. The flagellar oscillatory state was determined prior to each stimulation. **(C)** Normalized flagellar (displacement) and nerve (compound action potential) response envelopes are compared across conditions; oscillatory states: quiescent (left, *N* = 1), weakly SSO-ing (middle, *N* = 5), strongly SSO-ing (right, N=2); stimulation: one-tone (top), two-tone (bottom). The male flagellar sensitivity peak matches the female flight tone distribution (yellow highlight) across all conditions. The main sensitivity regions of flagellar and nerve responses, are mismatched during one-tone stimulation (top, blue/gray) across all oscillatory states, but match during two tone stimulation (bottom, red/black). A novel, large-amplitude nerve response emerges at higher frequencies (around 800 Hz, bottom left) despite minimal corresponding flagellar motion which gradually vanishes with increased SSO amplitude.

In addition to those stimulus conditions, male flagellar ears can exist in one of two qualitative mechanical oscillatory states: quiescent (non-oscillatory) or those exhibiting self-sustained oscillations, with the latter occurring at varying amplitudes. Based on the presence and strength of SSOs, animals were classified into three mechanically distinct states: quiescent, weak SSO and SSO (see Methods). Animals exhibiting SSOs have flagellar oscillation amplitudes exceeding 200 nm in the absence of external stimulation, whereas quiescent ones would have oscillation amplitudes of the order of 1 nm. Animals classified as having weak SSOs fell in the intermediate range between these two states. Together, this led to six mechanical input states (Fig.1C).

Across all six states, the envelope of the flagellar frequency response (grey curves in Fig. 1C) was broadly similar, resembling a forced damped harmonic oscillator with a resonance near the female wingbeat frequency (∼ 360 Hz), consistent with previous findings (*12*). Minor differences were observed in the SSO state under one-tone stimulation, these included small amplitude increases of the flagellar envelope near the SSO frequency and a slight increase in tuning sharpness, both consistent with a simple superposition of tones (SSO + stimulus tone).

The neural response outputs, however, showed stark differences across the states. In quiescent ears, and under one-tone stimulation (Fig. 1C, top, left), auditory nerve responses spanned a sensitivity range from 100 to 400 Hz, in line with previous studies (*5*, *8*, *10*, *13*, *14*). Sensitivity could be further subdivided into a main region between 100 to 300 Hz and a second, minor sensitivity region between 300 to 400 Hz (corresponding to ecologically relevant female flight tones). Animals undergoing strong SSOs (Fig. 1C, top right), in contrast, showed a novel higher-frequency region of audibility between 400 to 600 Hz. A similar sensitivity region was reported for *Culex* mosquitoes (*8*, *14*), but its link to the flagellar state was not explored.

In the two-tone series, nerve responses changed dramatically. In quiescent ears (Fig. 1C, bottom, left), three distinct sensitivity peaks could now be distinguished. The first major peak occurred between 100 to 300 Hz, which was broadly similar, to the main sensitivity region seen under one-tone stimulation. A second, and largest, peak emerged around the frequency range of female flight tones (300–400 Hz). This peak remained the largest across all flagellar states (Fig. 1C, bottom, left to right). Under two-tone stimulation, a third, and most remarkable, sensitivity region, emerged at higher frequencies, between 700–1000 Hz. Such high-frequency sensitivity was not seen under one-tone stimulation (Fig. 1C, top). This two-tone high-frequency sensitivity showed strong dependency on the flagellar oscillatory state. It was most prominent in quiescent ears (Fig. 1C, bottom, left) and completely abolished under strong SSOs (Fig. 1C, bottom, right). The newly created high-frequency audibility likely arises from *two-tone distortions*, as previously suggested for a lower frequency range in *Culex* (*5*, *8*) and *Aedes* (*15*) mosquitoes. Intriguingly, the magnitude of the nerve’s CAP responses in the here reported high-frequency region (700-1000 Hz) was similar to that seen in the lower frequency regions (100-400 Hz), although flagellar displacements are substantially smaller at the higher stimulus frequencies.

These findings raise two possible (but not mutually exclusive), interpretations: either the two-tone stimuli recruit a previously uncharacterized neuronal population specifically responsive to complex tonal patterns reminiscent of the proposed combination-sensitivity reported for vertebrates (*16*), or the resulting nonlinear distortions usher higher frequencies back into the neurons’ fundamental operating range (100–400 Hz), effectively generating *de novo* high-frequency audibility. This phenomenon can be seen as a form of biological heterodyning which creates “sound from ultrasound”.

### Matching distortions in flagellar receiver and nerve

To explore the landscape of audibility, the spectral composition of the responses was analyzed by performing Fast Fourier Transforms (FFTs) on both flagellar and nerve signals across all stimulus frequencies. The spectral decompositions were compiled into three-dimensional mosaics, where each column represented the spectrum at a specific stimulus frequency (Fig. 2A-D). These mosaics revealed distinct DP families, manifesting as straight lines, which take the form *α f*_1_ + *β f*_2_ + *γ f*_*sso*_, where *α*, *β*, *γ* are integers, *f*_2_ and *f*_*sso*_ are fixed tones corresponding to the male flight tone and the male’s SSO frequency, while *f*_1_ is varied (see Supplementary Methods).

**Fig. 2.**
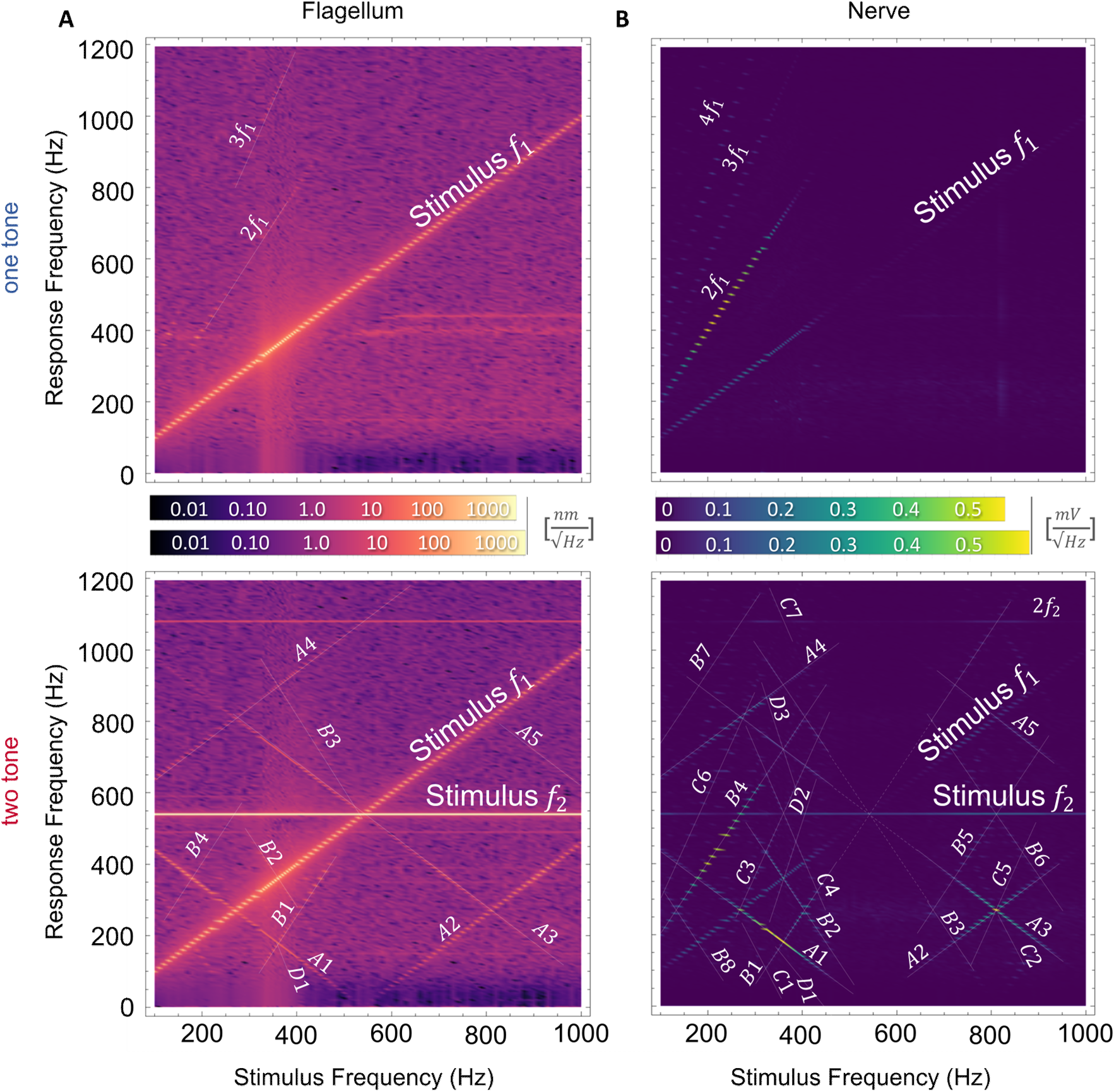
Spectral mosaics of flagellar and nerve responses in a quiescent animal. Distortion product families are labeled for identification (e.g., A1, A2, etc.) and marked with semi-opaque lines for clarity (see Supplementary Materials). **(A)** Spectral mosaics of flagellar displacements for one-tone (top) and two-tone (bottom) stimuli on logarithmic scale. Mosaics represent the FFT of the respective flagellar response, a first-order interpolation was applied for smoother visual transition between stimulus frequencies. Primary tones (*f*_1_, *f*_2_) dominate, harmonics and distortions are significantly smaller (10-100 times). Color legends reflect mechanical energies. **(B)** Corresponding mosaic of nerve responses to one-tone (top) and two-tone (bottom) stimuli on linear scale. For one-tone stimuli, the nerve is dominated by the frequency-doubled component reflecting the response from two anatomically opposing populations of neurons. Two-tone stimuli exhibit the two high-frequency response loci ( ∼300 Hz and ∼800 Hz), consistent with envelopes from Fig. 1C. The two-tone nerve response appears stronger than its one-tone counterpart, suggesting distinct neural dynamics under two-tone stimulation. Color legends reflect nerve response magnitudes. A list of all identified DPs can be found in table S1.

In quiescent animals, in the one-tone condition, the flagellar response was dominated by the primary stimulus tone *f*_1_ (Fig. 2A, top panel). While the primary tone was also present in the nerve spectrum (Fig. 2B, top panel), the nerve response was dominated by its second harmonic (2*f*_1_), a feature attributed to alternating firing of opposing neuronal populations in the Johnston’s Organ (JO) (*4*, *17*, *18*). Beyond 400 Hz, the neuronal response disappeared entirely, consistent with the known upper frequency limits of mosquito auditory neurons (*5*, *8*, *10*, *14*).

The two-tone condition (Fig. 2A, bottom panel) presented a more complex profile. In the flagellar response, the dominant components remained the primary tones *f*_1_ and *f*_2_, with additional contributions from quadratic DPs such as *A*1 = *f*_2_ − *f*_1_ and *A*2 = *f*_1_ − *f*_2_.

Notably, these quadratic DPs are distinct from beat tones, which represent amplitude modulations which are not spectrally visible. Furthermore, previously unreported cubic DPs, such as *A*3 = 2*f*_2_ − *f*_1_ and *B*1 = 2*f*_1_ − *f*_2_ were also detected. Despite these interactions, the total flagellar power increased only marginally due to the simple summation of the two primary tones.

In the nerve, however, the two-tone condition produced a far richer spectrum, with new DPs populating a wide range of frequencies, far exceeding that of biologically occurring pure tones. Not all nerve DPs could be traced to flagellar motion, suggesting contributions from other nonlinear elements within the nerve that are independent of the mechanotransducer complex attached to the flagellum. This is explored in more detail in the Discussion part of the Supplementary Information. The region spanning 100-400 Hz is densely populated with high-order distortions indicative of strong nonlinear processes being at play. The second region, between 400 and 600 Hz, was devoid of any nerve responses, including the primary tones, suggesting that male mosquitoes remain largely “deaf” to their own wingbeats even under two-tone conditions. The third region, from 600 to 1000 Hz, contained prominent components such the *A*2 quadratic, and the cubic *A*3 Given that these components were traceable to flagellar responses, they were likely signalled through mechanotransduction.

It is important to note that the auditory nerve itself does not show any direct sensitivity for (pure tone) frequencies above 400 Hz (Fig. 1C, top left). As it will be shown in shown in the following section, high-frequency combination tones are mapped onto lower frequency distortions which fall within the expected 100-400 Hz bandwidth of the nerve’s sensitivity range (Fig.4 C-E, and fig. S2). A more detailed discussion can be found in the Supplementary Text.

Control experiments showed that, once transduction is abolished using the chordotonal mechanotoxin pymetrozine (*19*, *20*), no distortions were observed - confirming their biological origin (fig. S1). These experiments also show that the presence of *f*_2_ in the nerve recordings is an artifact exclusively attributed to electrode crosstalk. There is no physiological response to *f*_2_which is in good agreement with a passive prefilter as previously suggested in vertebrates (*21*).

The auditory landscape of SSOing animals was dramatically different. In the one-tone condition (Fig. 3A), SSOs produced a constant, bright spectral stripe near 350 Hz, consistent with prior observations (*12*). A defining feature of SSOs is that they are able to entrain to external stimuli near their intrinsic frequency (in our case ∼350 Hz) (*11*, *22*, *23*). Entrainment, a phase-locking of the flagellar motion to an external stimulus, manifests as a “gap” in the SSO frequency around 350 Hz in which distortions are absent, consistently lowering the noise floor across all animals with an SSO amplitude larger than 50 nm.

**Fig. 3.**
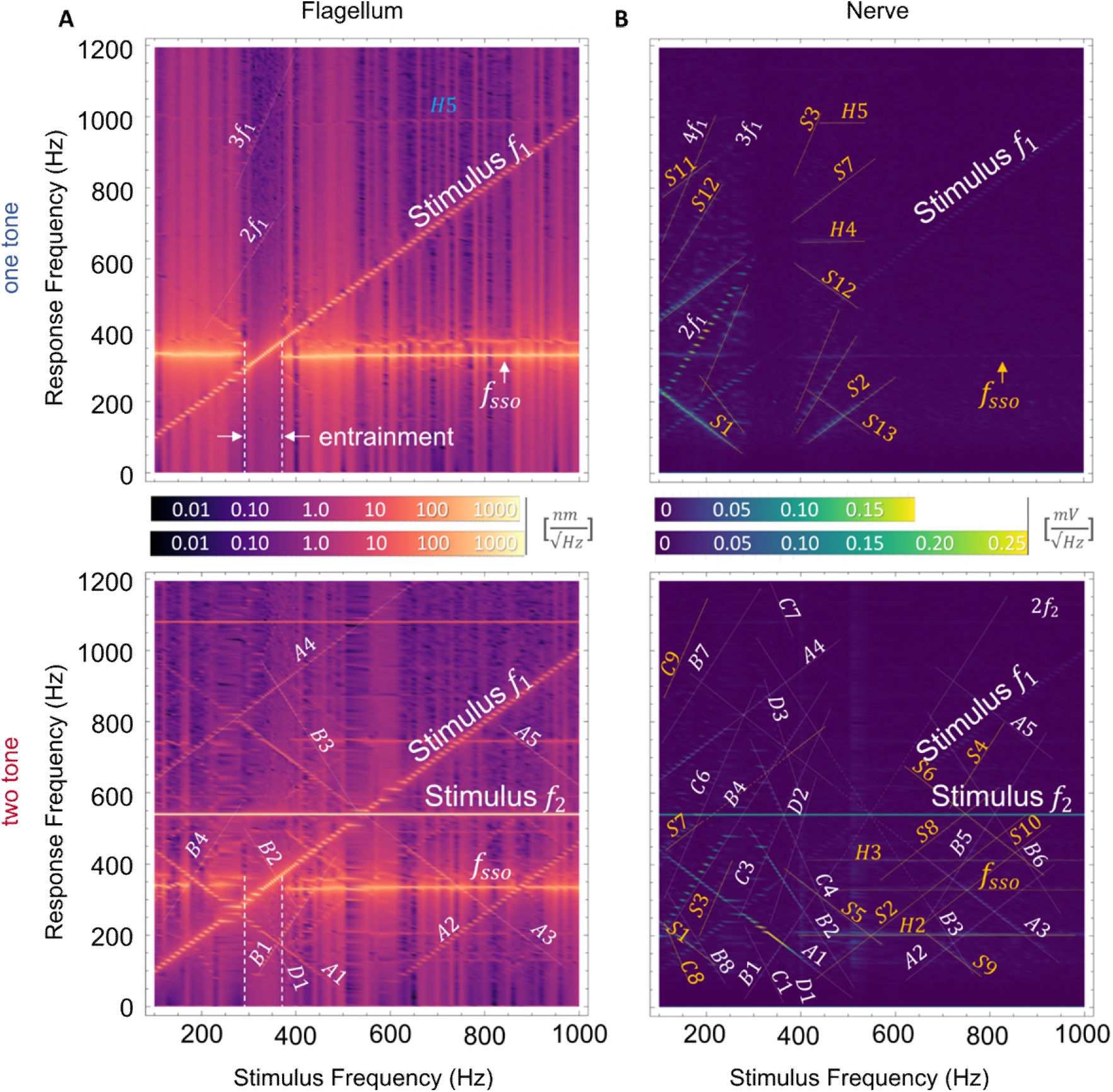
Spectral mosaics of flagellar and nerve responses in an animal exhibiting SSO. Distortion product families follow the same naming convention as in Fig. 2 but are highlighted in a different color to indicate SSO specificity. **(A)** Spectral mosaics of flagellar displacements. The SSO state manifests as a continuous band at *f*_*sso*_∼ 340 Hz. The band is broken upon entrainment (dashed vertical lines). In the two-tone case, the flagellum displays a greater number of distortions compared to the quiescent animal, albeit at low amplitude. **(B)** Corresponding mosaic of nerve responses. In contrast to quiescent animals (Fig. 2B, top), one-tone responses of SSO-ing animals show pronounced distortions, stemming from interactions with *f*_*sso*_. The entrainment region produces a zone free of distortions, suggesting that entrainment effectively dampens distortions at both the flagellar and nerve levels. In the nerve (bottom), the entrainment region persists, because *f*_*sso*_ phase-locks to the stimulus frequency. Other distortions persist in the same frequency region highlighting the complex interplay between SSO frequency and distortion damping, particularly in the context of two-tone stimulation. A list of all identified DPs can be found in table S2.

Nerve responses in the one-tone condition revealed that an entrained flagellar SSO silences all nerve responses indiscriminately, including the primary *f*_1_(Fig. 3B, top panel). Outside this region, multiple distortion products emerged due to complex interactions between *f*_1_and *f*_*sso*_, suggesting that the SSO state may have an active role beyond the previously suggested detection of conspecific females.

Under two-tone conditions (Fig. 3, bottom panels) both the flagellar and nerve mosaics of SSOing animals exhibited richer spectra than those of quiescent animals. Nevertheless, all observed distortions can be fully attributed to interactions among *f*_1_, *f*_2_ and *f*_*sso*_.

A striking observation was the substantial increase in total nerve output, with some distortion product families enjoying approximately 50% larger responses than those observed under one-tone stimulation. This suggests that nerve responses involving the *A*1 quadratic distortion may be able to recruit a larger number of neuronal units, pointing to heightened sensitivity to DP-based tones. The increased energy content observed in the SSOing two-tone condition likely reflects either tighter synchronization of neuronal firing or the activation of previously inaccessible neuronal populations.

These findings reveal that distortion products do not pollute the audibility landscape but subsume it.

### Neural sensitivity to distortions is enhanced

To break down compound nerve responses (Fig. 1C) into their individual constituent components, we extracted the neural response amplitude profiles for prominent DP families (Fig. 4A). In both quiescent (Fig. 4B,D) and SSO animals (Fig. 4C, E), compound nerve profiles could be reconstructed by accounting for a limited set of distortions. These included two quadratic distortions (*f*_2_ − *f*_1_ and *f*_1_ − *f*_2_), two cubic distortions (2*f*_1_ − *f*_2_and 2*f*_2_ − *f*_1_), and the nerve response to the primary tone (*f*_1_) along with its characteristic frequency doubling (2*f*_1_). In SSO animals, two additional quadratic distortions (*f*_1_ − *f*_*sso*_ and *f*_*sso*_ − *f*_1_) contributed to the one-tone profiles.

**Fig. 4.**
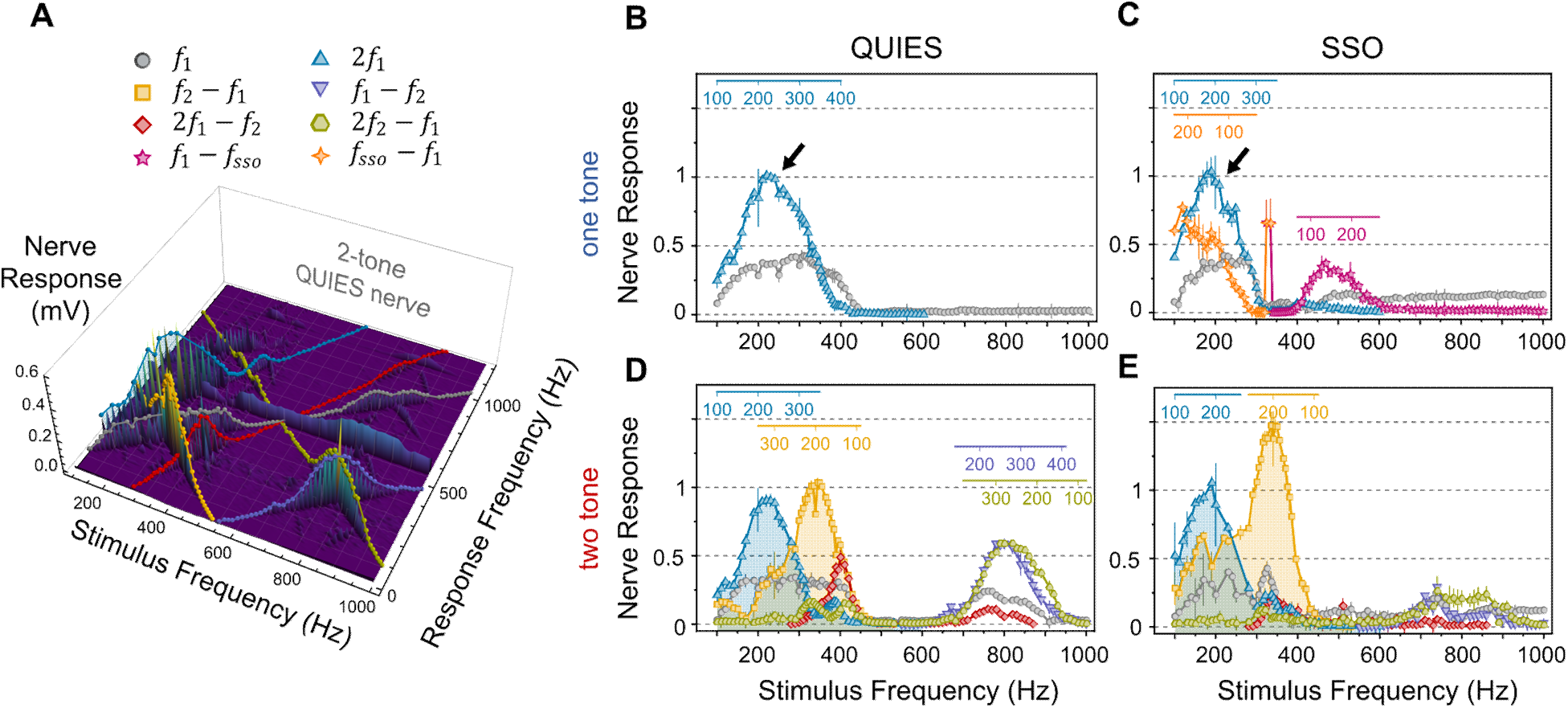
Nerve response profiles in quiescent and SSO-ing ears. **(A)** Amplitude profiles of prominent distortion products (DPs) from Fig. 2D. Legend applies to all panels. **(B-E)**, DP families are compared across quiescent and SSO conditions. The nerve response data for each mechanical state is normalized to its respective one-tone 2*f*_1_family (light blue, indicated by a black arrow). One-tone stimulation in quiescence (B), produces two response families. Both represent direct responses to the primary tone, which can manifest as single-frequency (*f*_1_, blue) or frequency-doubled response (2*f*_1_, light blue). Playing two tones instead (D) produce multiple DP response families in addition to the original ones. The high-frequency peaks mostly derive from two DP families: the quadratic *f*_1_ − *f*_2_(purple) and the cubic 2*f*_2_ − *f*_1_(green). (C, E) Amplitude profiles of DP families from SSO-ing ears (*N* = 3). The peak at ∼500 Hz in (C) corresponds to an interaction between *f*_1_ and *f*_*sso*_. In the two-tone case (E), the quadratic DP family shows significant amplification, likely due to flagellar SSO entrainment. It is noted that just a handful of DP families can fully explain the features in the response envelopes of Fig. 1C.

During two-tone stimulation in quiescent animals, the quadratic-cubic distortion pair of of *f*_2_ − *f*_1_and 2*f*_1_ − *f*_2_ contributes to the peaks observed in the low frequency region of the neural profiles (100 – 500 Hz; Fig. 1; Fig. 4D). In contrast, the quadratic-cubic pair of *f*_1_ − *f*_2_ and 2*f*_2_ − *f*_1_ accounts for the peak in the high-frequency region (600 – 1000 Hz; Fig. 1; Fig. 4D). In two-tone stimulated SSO animals, contributions in the high-frequency region appear suppressed, whereas a strong quadratic (*f*_2_ − *f*_1_) response – showing a ∼50% increase compared to the quiescent state characterizes the low-frequency region (Fig. 4E).

This increase in amplitude of the nerve response can be attributed to some combination of tighter neuronal synchronisation or the presence of a novel, combination-sensitive neuronal population. If the system relied exclusively on tighter neuronal synchronisation, the area under each DP amplitude trace would remain the same (Fig.4 and Supplementary Text). By calculating the ratio of areas between the quadratic response (*f*_2_ − *f*_1_, yellow) and the frequency doubled stimulus (2*f*_1_blue), one can estimate the “volume” of the neuronal pool that each distortion can use, relative to the fundamental one-tone neuronal population. In the quiescent case, the quadratic appears to have an 11% increase relative to 2*f*_1_ (Fig. 4D), whereas for SSOing animals the neuronal pool is measured to be 62% larger than the pure-tone counterpart (Fig. 4E). This is suggestive of the presence of a neuronal population that has specific, DP-based sensitivities which is more strongly recruited under SSO conditions.

In the one-tone stimulated SSO animals, the DP trace maxima for *f*_1_ − *f*_*sso*_ and *f*_*sso*_ − *f*_1_ depended on *f*_*sso*_, which varied across animals (fig. S4). Nonetheless, these analyses strongly suggest that previously reported high-frequency sensitivities of hearing in male mosquitoes (*8*, *14*) are directly linked to the presence of *f*_*sso*_ and its relation to the distortion *f*_1_ − *f*_*sso*_ (Fig. 4C).

Importantly, regardless of flagellar state, DP response frequencies consistently remained between 100 and 400 Hz, as shown by the colored inset axes (Fig. 4B-E), matching the baseline response range of the quiescent ear under one-tone stimulation (Fig. 4B).

The significant contributions of distortion products to mosquito neural responses presents an apparent disparity between the mechanics of the flagellar response and neural output. Similar to distortion product otoacoustic emissions observed in mammals (*24*), the magnitude of these distortions – as manifested in the vibrations of mosquito flagella – is quite small, typically 2-3 orders of magnitude smaller than the primary tones *f*_1_and *f*_2_(Figures 2, 3). Yet, despite their miniscule amplitude, these tiny distortions generate remarkably strong nerve responses in the mosquito ear.

To investigate this discrepancy, we examined the relationship between flagellar displacement amplitude and nerve response (Fig. 5). Contour plots were constructed for selected DP families (*f*_2_ − *f*_1_, *f*_1_ − *f*_2_, 2*f*_1_ − *f*_2_, and 2*f*_2_ − *f*_1_), as well as for responses to pure tones (*f*_1_ and *f*_1_, *f*_2_), mapping nerve response frequency against flagellar displacement.

**Fig. 5.**
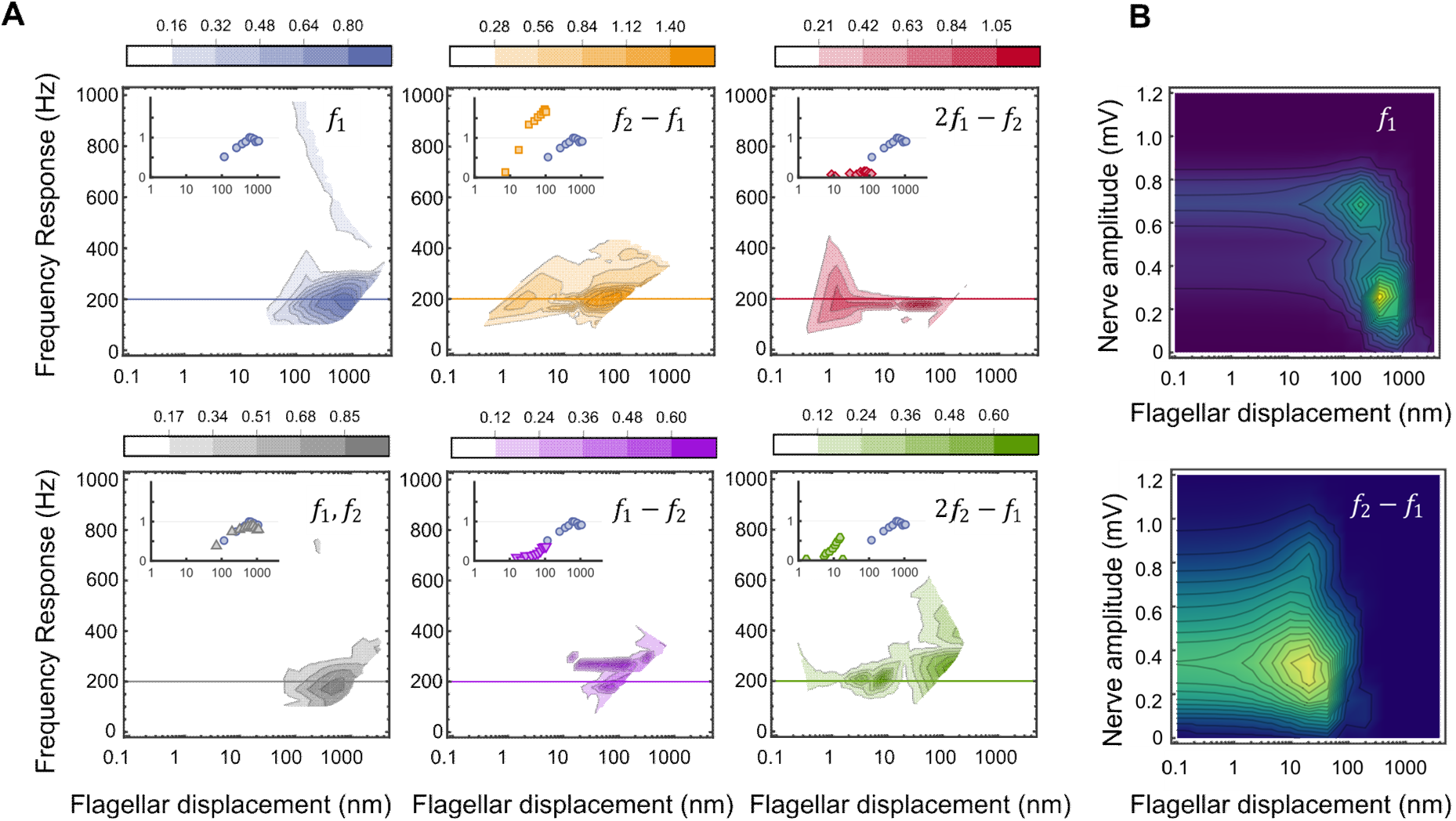
Topology of nerve responses as a function of corresponding flagellar displacement. **(A)** Contour plots of the relationship between flagellar displacement and nerve responses for the prominent DP families in Fig. 3E. The nerve response magnitude is normalized to the maximum of the *f*_1_topological map (in blue, corresponding to one-tone stimulus). Insets at top left of each plot illustrate a cross-section through the 200 Hz frequency response (highlighted horizontal line) with the corresponding normalized nerve amplitude on the y-axis. The panel labelled *f*_1_, *f*_2_ measures the response to *f*_1_ in the two-tone scenario. Topological maps show complexity of nerve responses and their dependencies on flagellar amplitude, frequency and stimulus type (primary tone or DP). Distortions require at least ∼10 times smaller flagellar displacements to generate a comparable nerve response. **(B)** Probability density histograms for the primary tone *f*_1_ (top), and the quadratic *f*_2_ − *f*_1_DP (bottom) across all studied animals independent of mechanical state (*N* = 11). The relationship between flagellar response and nerve response shows that the quadratic DP is universally more sensitive and elicits at least as powerful CAPs as via a pure tone equivalent stimulus frequency.

Notably, mosquito auditory neurons showed increased sensitivity for distortion products, requiring at least tenfold smaller flagellar displacements than pure-tone stimuli of the same frequency to generate comparable nerve responses (Fig. 5A). To further quantify this effect, we pooled maximal nerve responses and their corresponding flagellar displacements across all animals (regardless of their mechanical state), comparing the joint distributions of responses to *f*_1_and *f*_2_ − *f*_1_ (Fig. 5B). Median displacements were 695 nm for *f*_1_, but just 43 nm for *f*_2_ − *f*_1_, despite median nerve responses being of comparable amplitude (0.24 mV and 0.31 mV, respectively). A blocked PERMANOVA test comparing the centroids of the two distributions confirmed a significant difference (F(1) = 14.04, p-value < 0.001).

Discrepancies between the magnitude of distortion product otoacoustic emissions and their corresponding neural responses – where strong neural activity corresponds to a weak emission – have also been reported in mammals (*25*).

The striking sensitivity to distortions observed here challenges this canonical view of auditory, and possibly more generally neuronal processing. Using the mosquito as an exemplary case, our findings suggest that nonlinear interactions play a fundamental role in shaping neuronal signals and encoding sensory information.

### Low-frequency DPs retain high-frequency precision

To investigate the temporal properties of different phase-locked neural responses, we quantified the timing precision of compound action potential streams from their respective inter-peak intervals (IPIs; see Fig. 6A).

**Fig. 6.**
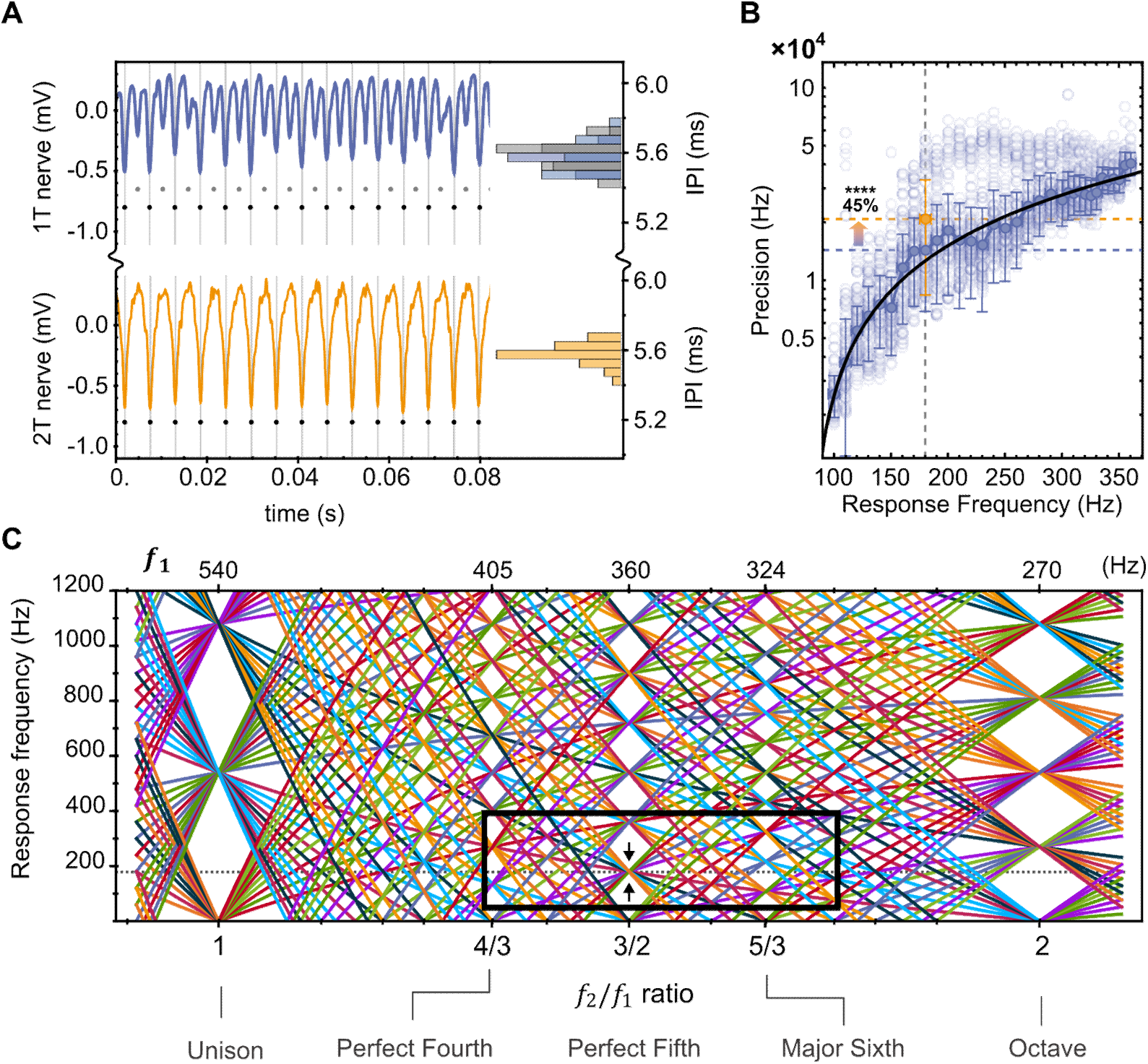
Precision and periodicity gains of distortion-based signaling. **(A)** Comparison of phase-locked neural responses (each at 180 Hz) from one animal. The top trace is a response from a pure 180 Hz tone, yielding two alternating streams from anatomically opposing populations of neurons (black and gray dots). In the bottom trace, a two-tone stimulus at a 3/2 ratio (*f*_1_ = 360 Hz) produces a single, sharply defined CAP stream, as multiple DP families converge (see panel C). Temporal precision is calculated from inter-spike interval distributions, shown as histograms beside each trace. **(B)** CAP precision from pure-tone (blue) and two-tone (orange) streams across all animals (*N* = 11). Black line shows linear fit to pure-tone data (log10 scale). At 3/2 stimulus ratios the temporal precision of responses is boosted by 45% relative to pure tones. **(C)** Simulated DPs across two-tone input ratios reveal spectral convergence at known musical intervals. Each colored line represents a unique DP family. Upper tick-marks indicate the corresponding *f*_1_for a given ratio. The 3/2 ratio funnels DPs into the *f*_1_ subharmonic (180 Hz, black arrows). The black box spans the ecologically relevant window for *Anopheles* mosquitoes, bound by the females’ WBFs (300-400 Hz, ∼19°*C*) and the males’ auditory range (100-400 Hz).

Precision data were extracted from all animals responding to one-tone stimuli with frequencies between 100 and 360 Hz, and for two-tone stimuli with *f*_1_at 360 Hz and corresponding distortions at 180 Hz (Fig. 6B). We compared the precision of direct responses to a 180 Hz one-tone stimulus with that of distortions at 180 Hz and found that distortions provided a boost in temporal precision of 45% relative to pure tones. A complementary Linear Mixed Effects Model analysis further revealed that the response type (a response to a one-tone vs to a distortion product) was a good predictor of temporal precision. Specifically, nerves producing a 180 Hz response via a distortion product exhibit greater temporal precision than through a pure-tone equivalent (Estimate=0.093, SE=0.024, t=3.80, p-value<0.001)

This finding is not trivial. It is a general truism of neural signaling that higher spiking frequencies confer greater temporal precision (*26*, *27*), while lower spiking frequencies are more energetically favourable (*28*, *29*) and less limited by the physical properties of neurons such as refractoriness (*30*). Here, distortions appear to offer the best of both worlds: a 180 Hz distortion in the male mosquito ear has half the frequency of a female’s flight tone yet recovers more than 50% of temporal precision in its phase-locked response (Fig. 6B).

Our mosquito analysis focused on two-tone distortions obtained specifically for *f*_1_ = 360 Hz, where the temporal component of the streams can be reliably extracted for the calculation of stream precisions (fig. S6). This limitation highlights a broader connection between distortion products and hearing. With an *f*_1_ at 360 Hz and an *f*_2_ at 540 Hz, the *f*_2_/*f*_1_ ratio is 3:2 (or 1.5), and the most dominant audible distortions produced in the mosquito ear will have a frequency of 180 Hz. Mosquitoes naturally have wingbeat frequencies such that the male-to-female frequency ratio is near 1.5 (*5*, *7*, *10*, *31*). For mosquitoes this wingbeat frequency ratio has been referred to as a “harmonic convergence ratio” or a “golden ratio” – in human terms, it is known as a musical ratio.

In music theory, a pair of tones with a ratio of 1.5 is called the “perfect fifth” - a musical interval of 3:2 - and is one of the most consonant and stable intervals in Western music. We thus wanted to use our mosquito model to examine more widely how the landscape of distortions varies across different tonal ratios, including classic musical intervals. To this end, we modelled all distortion products up to the 25th order (yielding a total of 598 DP families) as a function of the ratio between *f*_1_and the fixed male tone *f*_2_=540 Hz (Fig. 6C). Each line represents a unique DP family, with its frequency response plotted against the *f*_2_/*f*_1_ratio. Notably, for tonal combinations characterising musical intervals (unison, perfect fourth, perfect fifth, major sixth, octave), numerous distortions become mathematically identical, significantly reducing the distortional noise and complexity of the tonal landscape. This analysis also underscores that the only audible distortional frequency present when stimulating mosquitoes with *f*_1_ at 360 Hz and *f*_2_ at 540 Hz is 180 Hz.

Harmonically related tones are considered consonant because their waveforms periodically align, creating strong, easily identifiable (and harmonious) acoustic patterns. However, one might also speculate that, in a simplified scenario where the complexities and limits of pitch perception are relaxed (*32*), the harmony associated with these tones could be further enhanced by the relatively lower distortional noise generated within our ears.

## Discussion

From the opening of individual ion channels (*33*) to the operation of the entire brain (*34*), nonlinearities are integral to neural computation and have received particular attention in hearing research (*2*). A direct consequence of nonlinear filters is that they will invariably distort all signals passing through them. Whether such distortions are mere byproducts or vital elements of neural function has remained unclear. We used the flagellar ears of male mosquitoes to probe this question at the level of input-output dynamics across a range of tonal stimuli. We employed a dual approach, using analytically simple but ecologically relevant tonal stimuli. This allowed for a comprehensive analysis of mosquito hearing at its evolved optimalities. Our findings thus have implications for both neurobiology at large and mosquito auditory ecology in particular.

Our findings strongly suggest that essential distortions, which arise from the nonlinear mixing of two or more primary stimuli (e.g. pure tones) not only contribute to neural excitation but carry multiple benefits such as heightened firing regularity of neuronal units and an at least 10-fold increase in flagellar sensitivity.

For mosquito males chasing a single female within a swarm of thousands of other males, distortions provide audibility and corresponding spectral purity when the fundamental wingbeat frequency ratio reaches 3:2. In addition, the resulting distortion products are also isolated from the most salient frequencies of background noise. Most notably, the sets of audible distortions produced in the ear will depend on the chosen combination of (two or more) primary tones. Each pair of tones will generate a unique constellation of DPs, forming an acoustic fingerprint. The ability of the mosquito ear to distinguish pure tones from distortions of the same frequency will also help to distinguish ecologically relevant (flying conspecific females) from ecologically irrelevant stimuli (e.g. other insects with flight tones around the distortion frequency). Even if different tone pairs produce the same dominant response frequency, the overall nerve response remains distinct due to the unique pattern of DPs associated with each stimulus pair. Within a landscape of acoustic fingerprints, neuronal populations coupling to specific distortion frequencies would serve as a simple yet robust substrate for extracting complex acoustic features.

These findings also place the mosquito ear within a broader cross-species framework of auditory processing, highlighting surprising parallels with vertebrate hearing. Much like songbirds, bats, primates, and frogs, which use combination-sensitive neurons (*16*) to decode complex acoustic patterns, mosquitoes appear to employ a peripheral nonlinear strategy to enhance ecologically relevant signals. This suggests that the fundamental principles governing distortion-based hearing are likely shared across taxa, making mosquitoes a powerful model for studying nonlinear auditory mechanisms more widely. Their mechanically driven auditory processing could even offer a biologically accessible, invertebrate, *in-vivo* alternative to traditional vertebrate models to investigate distortion-based encoding in sensory systems.

DPs are as ubiquitous in the ears of vertebrates as reported here for mosquitoes. But due to the small magnitudes by which they manifest themselves, e.g. in recordings of otoacoustic emissions, they have been dismissed as proper stimuli in their own right. However, such easy dismissal could be premature: the heightened sensitivity to distortions observed for mosquitoes might also apply to vertebrates - the underlying mechanisms of which are poorly understood and merit deeper scientific scrutiny. Here, we offer a system where low-amplitude DPs are functionally superior to large amplitude primaries as the auditory system can fully exploit its nonlinear properties (*35*, *36*).

And finally, a distortion-based logic may hold value beyond auditory biology. By transforming high-frequency signals into metabolically cheaper, low-frequency representations with retained temporal fidelity, distortions could not only form a vital part of how neurons process signals and transmit information, but also a vital part of how we should stimulate neurons. Devices such as cochlear implants or new types of neural interfaces or sensory prosthetics might benefit from encoding principles that use structured nonlinearities - not despite their produced distortions, but because of them.

## Acknowledgments

We thank Dr Rainer Beutelmann for valuable discussions throughout the preparation of this manuscript.

## Author contributions

Conceptualization: AA, MG, JTA

Methodology: AA, MG, JB

Investigation: MG

Validation: MG, AA

Formal Analysis: AA, MG

Visualization: AA

Resources: JB, MG, AA

Funding acquisition: MA, DB, JTA

Project administration: JTA

Supervision: JTA, DB

Writing – original draft: AA, MG

Writing – review & editing: AA, MG, JF, JB, MA, DB, JTA

## Data and materials availability

The authors declare that the data supporting the findings of this study are available within the paper, its supplementary information files, and will be made publicly available on a public repository (i.e. Zenodo) upon publication of this work.

All analytical pipelines and plots in this manuscript were made using Wolfram Mathematica v.13.1 (https://www.wolfram.com/mathematica/), and open-access Python libraries. The authors will made the code accessible on a public GitHub repository (https://github.com/) upon publication.

## Supplementary Materials

### Materials and Methods

#### Mosquitoes

All experiments used 3–7-day-old male *Anopheles gambiae* (G3 strain). These were entrained in environmental incubators and reared under 12h:12h Light:Dark cycles (lights on at *Zeitgeber Time* ZT0), as described (*12*). Mosquitoes were removed from the incubators to be used in experiments around ZT12 (±3 hours).

#### Experimental preparation

Mechanical and electrophysiological measurements were obtained from mosquitoes in all experiments. Mosquitoes were mounted under the experimental apparatus following previously described procedures (*11*). Data were collected from 14 animals in total. All experiments were conducted at a room temperature of 18 − 19°*C*.

#### Stimulus Preparation

The mechanical motion of the flagellum and the respective nerve response at the JO are recorded simultaneously with a sampling resolution of 20 kHz as detailed in (*10*, *12*). The stimulus frequencies span between 100-1000 Hz in increments of 10 Hz by exception of the range between 300-400 Hz where the spacing is 5 Hz instead. This region is of particular interest as it reflects both the typical female WBF as well as the typical region of flagellar entrainment to an incoming stimulus.

The one-tone stimulus consists of a pure cosine wave of frequency *f*_1_, representing a flying mosquito’s – male or female – wingbeat frequency (WBF), the amplitude of which rises and falls (‘attack-and-decay’ ramp) over the course of 4 seconds to mimic its approach and departure. In the two-tone stimulus, a fixed second tone is superimposed at *f*_2_ = 540 Hz, corresponding to an average male mosquito’s typical WBF at 19 °C (*37*). The amplitude of the *f*_2_ stimulus was determined by measuring the typical wingbeat intensity of a tethered male mosquito at the level of the flagellum (represented as a command voltage of 2.5 V). To determine the maximum amplitude of the ramped tone, *f*_1_, several values were tested, and the highest value was chosen so that under single-tone stimulation, most male nerves only briefly reached saturation (maximum command voltage of 0.5 V). These procedures ensured that the stimuli remained ecologically relevant.

Each stimulus was divided into 20 segments with 10 amplitude divisions per slope, with segment 5 selected as representative for the main text analysis. Detailed amplitude dependency results are presented in the supplementary materials.

Each recording for a given stimulus (whether one-or two-tone) is screened for experimental errors based on the flagellar laser Doppler vibrometry (LDV) recording. If the recording signal drops below a specific threshold (*12*), the entire stimulus (flagellar and nerve) is discarded.

Discarded datapoints can manifest as “smears” on the spectral mosaics as a 1st order interpolation is used across all spectra for visual clarity.

#### Separating animals based on flagellar mechanical state

Each stimulus (whether one-tone or two-tone) was preceded by a silent period, allowing the flagellum to return to its relaxed state. This pre-stimulatory period was used to classify the flagellar mechanical state as either quiescent or SSOing (spontaneous self-oscillating) (Fig. 1B Classification was performed by analyzing the amplitude distribution (*12*) or by examining the FFT spectrum, as SSOs typically appear as a single frequency between 300 and 400 Hz.

As the SSO amplitude played a major modulatory role in the nerve’s auditory responses, animals were classified based on the characteristic DP families emerging in the nerve across the 100–1000 Hz frequency range. This classification was further validated using principal component analysis (PCA) applied to the flagellar envelope profiles. PCA revealed two primary clusters—quiescent and SSOing—along with an intermediate group, weak SSO, which exhibited the greatest variability in DP families (fig. S9). Specifically, quiescent animals were defined as those whose flagellum oscillated by less than 1 nm in the absence of stimulation, while SSOing animals exhibited a minimum oscillation amplitude of 200 nm within the 300–400 Hz frequency range. In contrast, weak SSO animals (1–200 nm oscillation amplitude) formed a continuum, transitioning smoothly from a quiescent to a fully SSOing state in terms of their resulting DP families.

When clustering was performed using nerve envelopes rather than flagellar envelopes, the resulting PCA revealed high-order component matrices, rendering them insufficient as high-dimensional components would be necessary to explain the system’s total variance. However, the fact that PCA applied to flagellar envelopes verified clustering animals based on their DP families indicates that flagellar mechanics serve as a strong predictor of the DP families observed in the nerve.

#### Distortion product identification and order

Identification is done at the level of flagellar and nerve mosaics, which are constructed for discrete amplitude segments during the stimulus ramp. Each column in the mosaics corresponds to an FFT spectrum of the system for a given stimulus frequency (either one-or two-tone).

Before identification, a peak detection algorithm is run on the FFT data of the flagellar and nerve responses. All extracted peaks are curated by passing them through fixed criteria on the size, sharpness, and survival after Gaussian blurring, to avoid identifying noisy peaks. The resulting “map” is a dimensionally reduced system of what is shown in Figures 2 and 3. A custom-made algorithm collates the FFT peaks and identifies the DP family lines. Each DP family must contain at least 5 prominent peaks across the entire stimulus range, and each one of them must be maximally 20 Hz apart from each other, to avoid the identification of sporadic, stochastically defined DP families.

All animals show small variations on the number of DP families or of their amplitude profiles. The inter-individual variation of flagellar resonance frequency, or even the difference in amplitude or frequency of SSOs can affect which distortions, and at which frequency regimes, will manifest. Despite these variations, distortion products follow mathematically robust rules where the emergent frequencies must be the sum, or difference, of any integer-multiple of the primary tones involved.

To account for all the variations that can affect the identification of a DP family across a spectral mosaic, a simple linear model is fit that takes the form of *α f*_1_ + *β f*_2_ + *γ f*_*sso*_, where the coefficients *α*, *β*, *γ* are integers. For quiescent animals which do not have a sufficiently strong SSO presence in the flagellar or nerve response, the model is reduced with *γ* = 0.

The useful metric of nonlinearity is the distortion product *order*, given by the sum of the coefficients. Distortions with large DP order typically indicate that stronger nonlinear effects take place. Nonlinear processes that strongly affect the primary tone *f*_1_ manifest in the spectral mosaics as lines with sharp slopes (both positive or negative ones). By contrast, nonlinear processes that affect *f*_2_or *f*_*sso*_ more strongly will lead to a stronger vertical shift of the entire DP family line in the mosaic.

A useful metric of nonlinearity is the distortion product order, given by the sum of the coefficients *Dp_order_* = |*α*| + |*β*| + |*γ*|. Distortions with large DP order typically indicate that stronger nonlinear effects take place. It is noted that typically higher order DP families are described by smaller response amplitudes with lower contributions to the total response output (*38*). Nonlinear processes that strongly affect the primary tone *f*_1_ manifest in the spectral mosaics as lines with sharp slopes (both positive or negative ones). By contrast, nonlinear processes that affect *f*_2_ or *f*_*sso*_ more strongly will lead to a stronger vertical shift of the entire DP family line in the mosaic as *f*_2_is fixed at 540 Hz by design, and *f*_*sso*_ is largely constant in frequency over the course of an entire experimental series recording.

#### Pymetrozine Injections

Pymetrozine is a neuroactive compound that acts as irreversible agonist (‘open channel blocker’) of the dimeric channel complex formed by the two transient receptor channels Nanchung and Inactive (*19*, *20*), which are important for neuronal signalling in insect chordotonal organs (*39*). Injection of pymetrozine has been demonstrated to abolish local field potential responses in mosquito ears (*11*). Consequently, it was here employed as a control to ascertain the biological origin of the observed distortions in mosquito auditory nerve recordings. The injections were conducted in accordance with previously described procedures (*11*, *40*).

#### Temporal precision analyses

Temporal precision analyses included comparisons between the IPIs extracted from the nerve responses to single tones with frequency *f*_1_ and nerve responses to distortion products with frequency *f*_1_. One difficulty with extracting temporal information of the neural recordings resided in the electrophysiological method used for measurements. Specifically, the local field potentials that were measured constitute the summed output of all the active neurons in the mosquito Johnston’s organ. During two-tone (or in the case of a non-entrained SSO animal, three-tone) stimulation, many of these neurons had different frequency responses and phase sensitivities, resulting in a complex waveform from which useful spike-related temporal information is difficult, and often impossible, to obtain (Fig. S6).

It was nonetheless possible to extract IPIs from the neural responses to single tone stimuli. It was also possible to extract this information from responses to two-tone stimuli when the ratio between the two stimulating frequencies was 1.5 – all distortions produced have the same frequency at that ratio. IPIs were extracted using a graph-based method: First, a peak detection algorithm was applied to neural traces, extracting peaks (inverted “spikes”) above a pre-defined noise threshold. The latter was computed from the nerve recording over the period of silence prior to stimulus presentation and thus varied for each extraction. An independent noise threshold was computed for each stimulation. Pairs of peaks were then connected if the instantaneous frequency (reciprocal of IPI) between them was within 15 Hz of the firing frequency of interest – the 15 Hz margin was found to strike a balance between peak finding and computational costs. These peak-to-peak connections were then used to form a graph where each peak constituted a node. The graph was traversed to extract series of peaks, or “streams of firings” reflecting a set of neurons responding with the given frequency of interest.

Temporal precision was computed from the IPIs of extracted streams. A jitter-derived metric was estimated as half the standard deviation of the IPIs computed from a stream.

Precision was calculated as the reciprocal of this jitter value.

#### Linear Mixed Effects Model Development

A linear mixed effects model (LMEM) was employed to analyse temporal precision extracted from neural streams across all animals regardless of mechanical state. Specifically, the analysis compared responses elicited by one-tone stimuli (*f*_1_ at 180 Hz) with those obtained from two-tone stimulation (*f*_1_at 360 Hz and *f*_2_at 540 Hz) that generated distortions at 180 Hz.

Temporal precision, the outcome variable, was *Log*_10_-transformed to satisfy normality assumptions. The sole fixed effect was the categorical predictor ‘response type’, with two levels: ‘one-tone at 180 Hz’ and ‘distortions at 180 Hz’. To account for individual differences, mosquito identity was included as a random intercept. The LMEM was applied to temporal precision estimates derived from the first five segments of the stimulus ramp (see “Methods - Stimulus Preparation” for more information).

#### Limitations

The mechanical state of an animal (SSO or QUIES) exhibits variations over the course of the entire experimental series, meaning that both mechanical properties of the flagellum as well as the amplitude profile of DPs can vary over time for the same animal. Although the SSO frequency remains largely constant (*σ* = 5 Hz) and leads to only minor vertical shifts of DP traces on the spectral mosaics (Figures 2 and 3), the amplitude fluctuations are more varied and can reach up to 25% of the average oscillation amplitude in size. The effect or role of those fluctuations are more difficult to quantify. Despite that, no animals exhibited fluctuations strong enough to push them out of their designated mechanical state classification (quiescent, weak SSO, SSO).

#### Cross talk analysis

Presentation of auditory stimuli via electrostatic actuation sometimes resulted in cross-talk between the neural recordings and the stimulus, with the stimulus being picked up by the nerve’s recording electrode. This was generally more pronounced for the constant, high-amplitude tone *f*_2_. To remove *f*_2_-related cross-talk, all two-tone stimuli include 100 ms at the start and end which only contain *f*_2_. These sections were used to fit a sinusoid to the corresponding nerve segments. This fitted sinusoid was then subtracted from the nerve response, yielding a signal free from *f*_2_-related cross-talk. The associated *f*_2_cross-talk is deliberately left in Figures 2 and 3 as a visual aid.

Force step stimulation experiments were used to assess cross-talk associated with *f*_1_. These experiments were conducted as described previously (*11*, *41*). For each step size used, averaging over several stimulus presentations allowed for removal of transient cross-talk and enabled assessment of compound action potential (CAP) amplitudes in response to the step. The CAP amplitudes were determined by applying a peak detection algorithm to the averaged trace. By taking the absolute value of each response, averaging across all presentations for each step size, and applying the peak detection algorithm to the first millisecond of stimulation, the amplitude of the electrostatic actuation cross-talk picked up by the recording electrode was computed. A cross-talk ratio, defined as the ratio between CAP amplitude and the cross-talk amplitude, and was then used to assess the relative level of cross-talk exhibited by each animal. Animals exhibiting cross-talk ratios greater than 0.2 at command voltages similar to the highest command voltage of stimulus *f*_1_ were discarded from subsequent analyses of temporal precision (see “Stimulus Preparation” in the main text Methods).

### Supplementary Text

#### Nerve distortion products with no apparent flagellar equivalent

In the one-tone case of SSOing animals (Figure 3 in the main text), the distortions S1 and S2 have a strong prominence in the nerve output but do not appear to have an equivalent one in the flagellum (shown in the logarithmic scale). This is a rather unique case as in quiescent animals all major DP families have a flagellar analogue which points at the flagellum as a potential source of nonlinearity that the JO can process.

The lack of SSO-specific distortions in the flagellum could be attributed to one of three possible scenarios: 1) the nonlinearities that generate these distortions occur either at, or after mechanotransduction, 2) limitations on adequately recording the flagellar motion, or 3) selective JO filtering where specific distortions are enhanced in the nerve but suppressed in the flagellum through active or passive mechanisms.

The simplest scenario is one where the resulting S1 and S2 distortions occur either at, or after mechanotransduction. Pharmacological interventions with pymetrozine (fig. S1) show that although all distortions in the nerve are abolished, those in the flagellum remain largely intact and resemble those seen in purely quiescent animals. Pymetrozine is not a transduction blocker *per se*, but rather acts as irreversible (open-channel) blocker - or agonist - of a dimeric transient receptor potential channel (Nan/Iav), which is deemed to be downstream of the actual transducers and required for the normal transformation of transducer currents into action potential responses (*20*). However, previous work on pymetrozine injections in mosquitoes has shown that it also affects flagellar stiffness, indicating that the JO can exert a certain degree of influence on receiver mechanics (*42*). The fact that certain prominent DP families retain their form in the flagellum even after pymetrozine injections point towards the fact that these nonlinearities originate, at least partially, at the flagellar level. Conversely, DP families which are present in the nerve but absent in the flagellum could indicate at a nonlinearity at the mechanotransduction level as previously proposed by the gating-spring model (*23*, *41*). Such a system supports the idea of “combination-sensitive neurons”, which respond preferentially, or uniquely to certain distortion products. Such DP-specific neurons have been observed in bats, monkeys, songbirds and frogs (*16*).

Another plausible mechanism would be due to resolution limitations of the laser Doppler vibrometer. Although the current threshold imposed in the mosaics is set to ∼1 nm, it is possible that the mechanical distortions are below this threshold. Further experiments where the dynamic range of the LDV is set closer to low-amplitude stimuli could clarity this possibility. It is noted however that for small stimuli, the flagellar stiffness measures roughly 150 *μN*/*m* (*11*) which would yield a thermal background at 7 nm at 20 °*C*, well above the LDV’s recording resolution indicating that it is unlikely that the recordings are resolution-limited. On the other hand, it is quite likely that distortions may be spatially localised. As the flagellum is not a uniform structure and a single laser measurement point may not be reliable it is possible that certain distortion frequencies may have an oscillation node at the point of measurement rendering them invisible to our recordings. The usage of scanning-, or two-axis vibrometry could rule out such possibilities. Lastly, the DPs observed at the flagellar and nerve level could be in part originating at the JO. Both fruit flies and mosquitoes have been shown to actively modulate their mechanical sensitivities to incoming sound (*11*, *43*) pointing towards an active intermediary process introducing nonlinearities. The presence of SSOs already suggests the capacity for an internal force generation even though the flagellum is significantly larger organ than auditory hair cells in vertebrates. It is thus reasonable that some emergent distortions are not due to external stimuli alone. To that extent, it is possible that internally generated distortions are not symmetrically presented to the flagellum and nerve leading to a mismatch between the two.

#### Frequency mismatch between flagellar and nerve responses

The range of nerve responses during two-tone stimulation is, at first glance, surprising (Fig. 2B, bottom; Fig. 3B, bottom), since it was shown that the frequency bandwidth to which the mosquito auditory nerve is sensitive (able to respond with phase-by-phase compound action potentials) to is limited to approximately 100 – 400 Hz (Fig. 1C, top left). However, distortional responses apparently extend beyond this range, reaching frequencies higher than 400 Hz (Fig. 2D). This discrepancy can be explained by considering the nature of the recorded signal, and inherent interpretational limitations of Fourier transform analysis in studying nerve responses.

Since the measured quantity is a local field potential, it represents a summed response of multiple active neurons during stimulation. This resulting complex waveform includes contributions from distinct neuronal populations firing at various frequencies, potentially overlapping in the recorded signal (fig. S3). This complex waveform may encompass not only compound action potential responses but also receptor potentials capable of tracking frequencies higher than 400 Hz – but do not generate spikes and do not propagate beyond their area of origin. Indeed, research on *Drosophila* indicates that local field potentials reflect ion channel open probabilities, suggesting they capture underlying receptor currents and potentials, potentially revealing high-frequency spectral components (*17*).

Other higher frequency components in the responses can be understood differently. As the FFT produces peaks only for reliably periodic signals, the presence of high frequency components could naturally emerge if certain neuronal units fire at a fixed lag or advance with respect to the majority of the responding neurons, in effect producing a secondary spike which accompanies the “main” neuronal spike. If this temporal “offset” between this secondary and main spike is reasonably fixed but small, it would manifest in the FFT spectrum as a high frequency component. Nevertheless, as is shown (Fig. 4 of the main text and fig. S2), the frequency space occupied by the dominant DP families maps nicely to the 100 – 400 Hz bandwidth of the nerve’s sensitivity.

**fig. S1.**
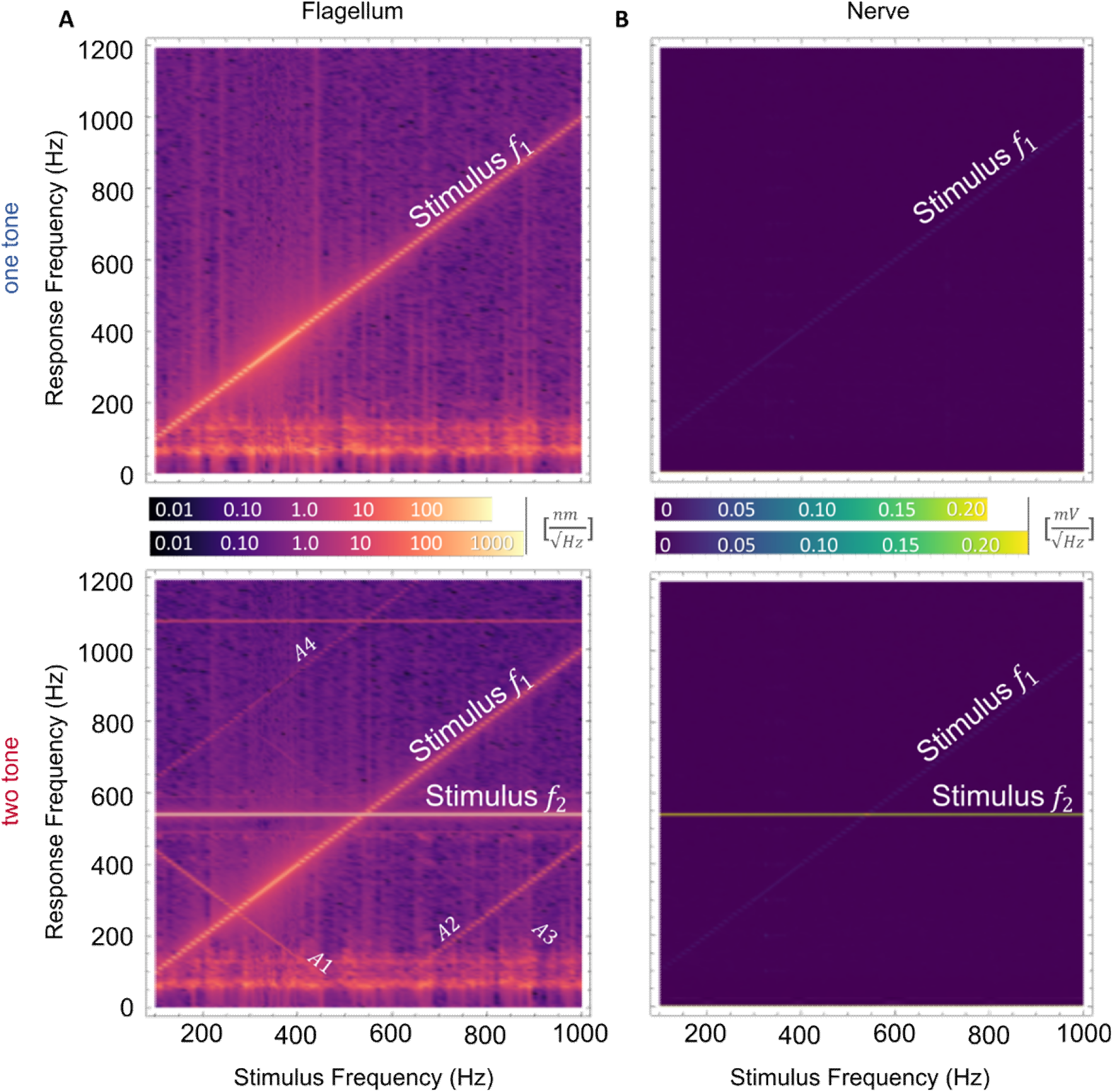
Spectral mosaics of flagellar and nerve responses after pymetrozine injections. **(A)** Spectral mosaic of the flagellar to a one-(top) and two-tone stimulus (bottom). Although the distortions present here largely resemble those seen in quiescent animals (Fig.2A) they distinctly lack the harmonics of the *f*_1_stimulus pointing towards a coupling between the generation of harmonics and mechanotransduction. **(B)** Spectral mosaic of the nerve response to a one-(top) and two-tone stimulus (bottom) under pymetrozine conditions. All nerve responses and distortions are completely abolished leaving behind a trace due to cross-talk between the stimulation and recording electrodes.

**fig. S2.**
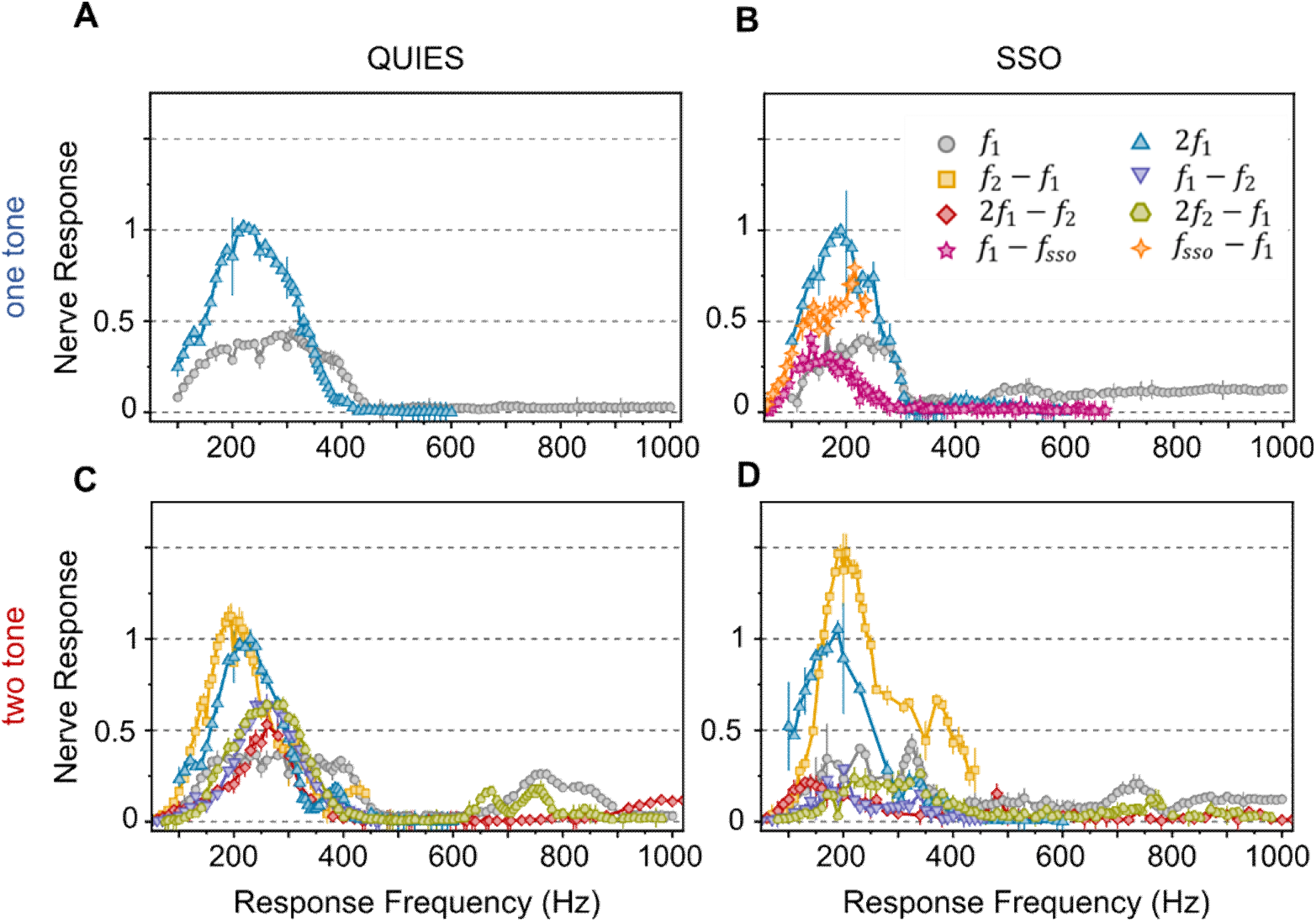
Nerve response traces for quiescent and SSOing animals as a function of the response frequency. These plots use the same DP traces shown in Fig.4B-E with the x-axis is recast on the nerve’s response frequency rather than the stimulus frequency. The y-axis represents the normalized voltage response of the nerve recordings. All distortion product families show to respond between 100-400 Hz meaning that DPs exploit the same fundamental neuronal architecture as their pure-tone counterparts. The **2*f***_1_ trace (light blue) has been manually remapped by halving the response range to account for the two opposing neuronal populations.

**fig. S3.**
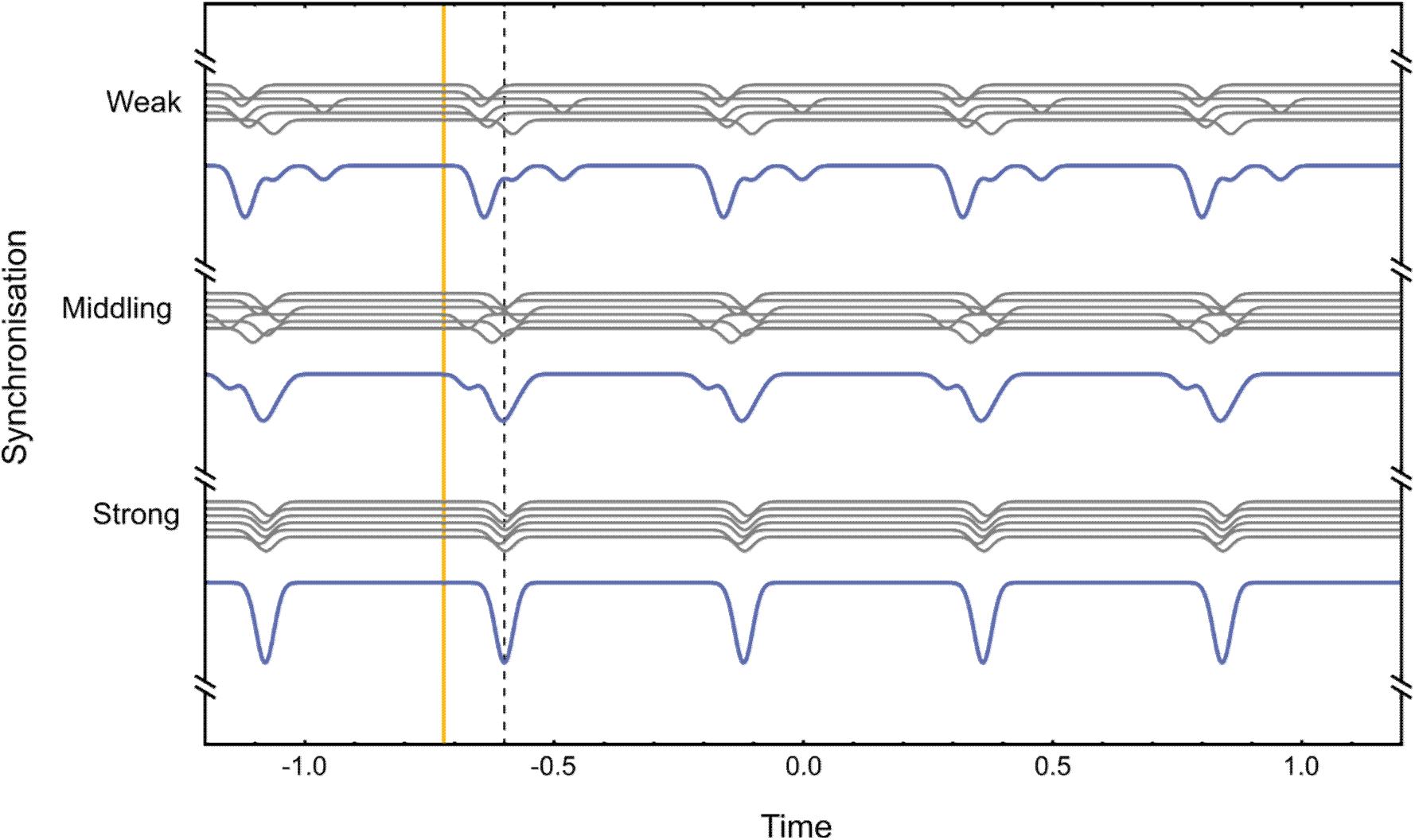
Tighter neuronal synchronisation leads to sharper, well-defined CAPs. Simulation of a nerve (blue) comprised of 6 neurons (gray). Each neuron is modelled to be perfectly precise and reliable, but the degree of neuronal synchronisation is varied across each panel (weak, middling and strong synchronisation). The orange and gray gridlines offer a visual reference on the neuronal synchronisation across each panel. While the relative phase between neurons is large, the resulting CAP is poorly defined and temporally spread out. Tighter synchronisation will not only yield stronger nerve responses, but the resulting spectra would also be less complex.

**fig. S4.**
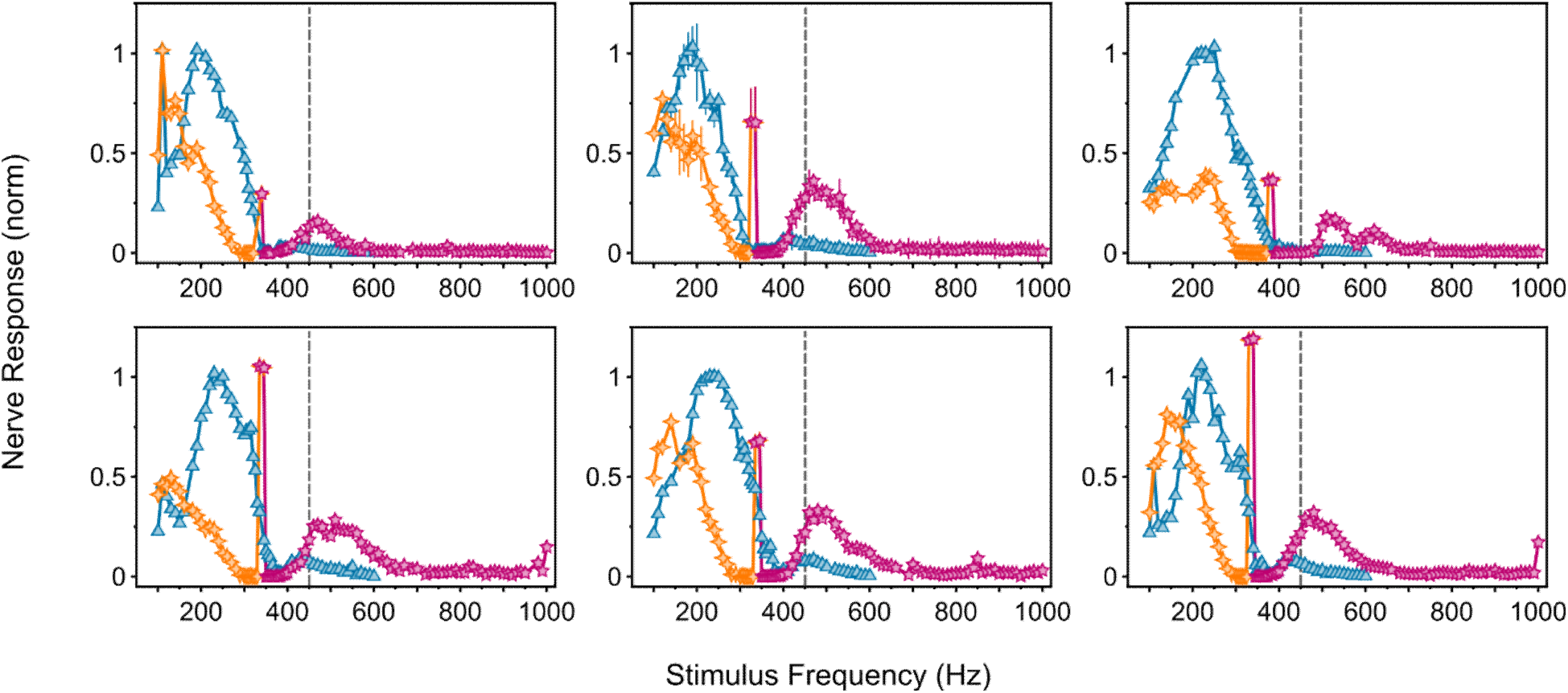
Variability on SSO-related DP families, based on the characteristic *f*_*sso*_. The DP traces to one-tone stimuli of various SSOing animals of various strengths is shown. Of particular prominence are the ***S*1** (orange) and ***S*2** (pink) DP families, whose size and relative position depend on the intrinsic SSO amplitude and frequency. The size of entrainment is also indirectly shown by the connecting regimes between ***S*1** and ***S*2**. The centerpoint of the two always matches the fundamental SSO frequency of each animal, occurring at the ***S*1** = ***S*2** intersection. The dashed line is set at 450 Hz as a visual reference.

**fig. S5.**
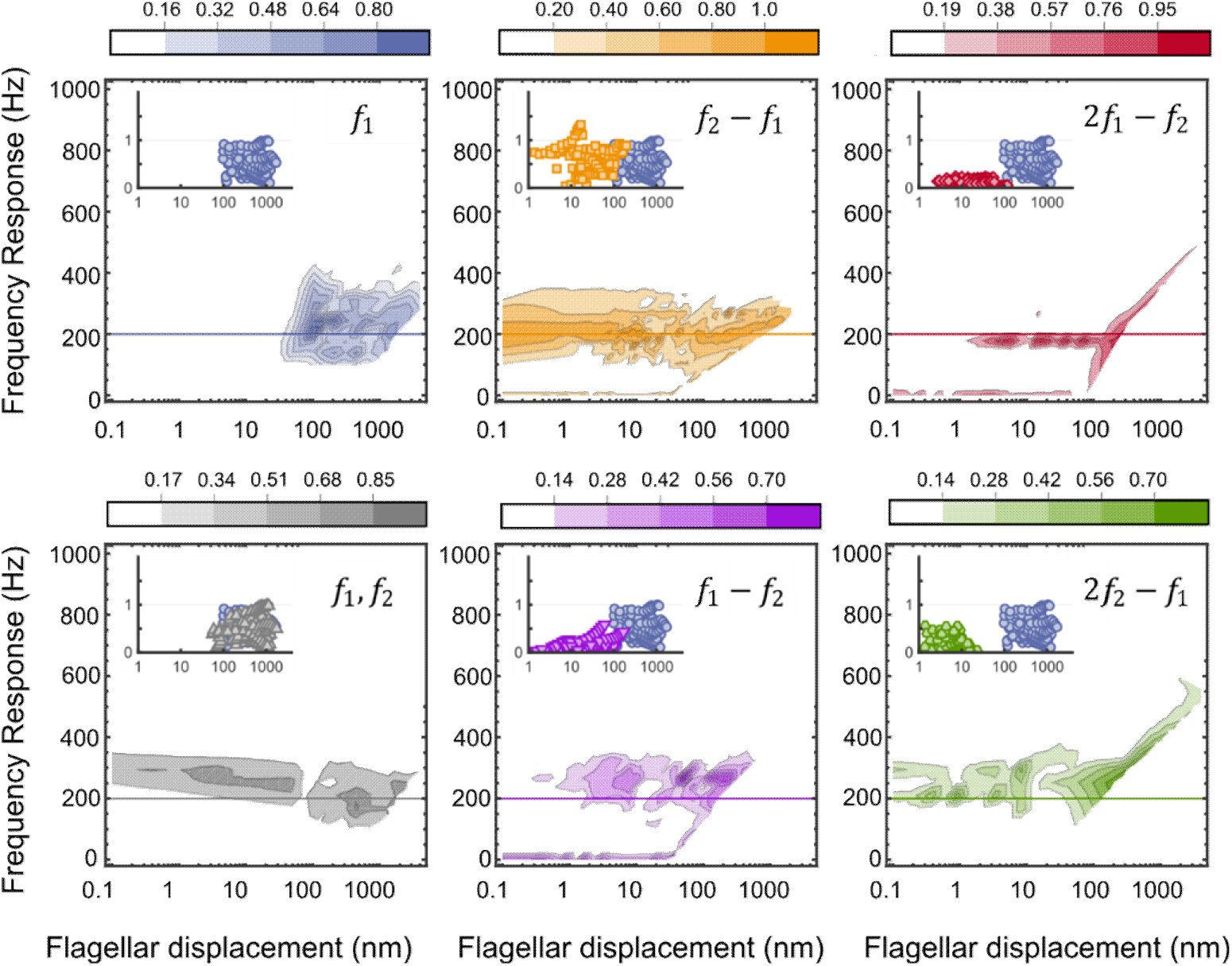
Topology of nerve responses as a function of corresponding flagellar displacement. Contour plots of the relationship between flagellar displacement and nerve responses for some prominent DP families across all animals regardless of their flagellar mechanical state. Similarly to Fig. 5a, nerves respond to stimulation with frequencies that lie between 100-400 Hz. Although different animals can have qualitatively different response topologies, DPs outperform pure-tone responses by at least one order of magnitude.

**fig. S6.**
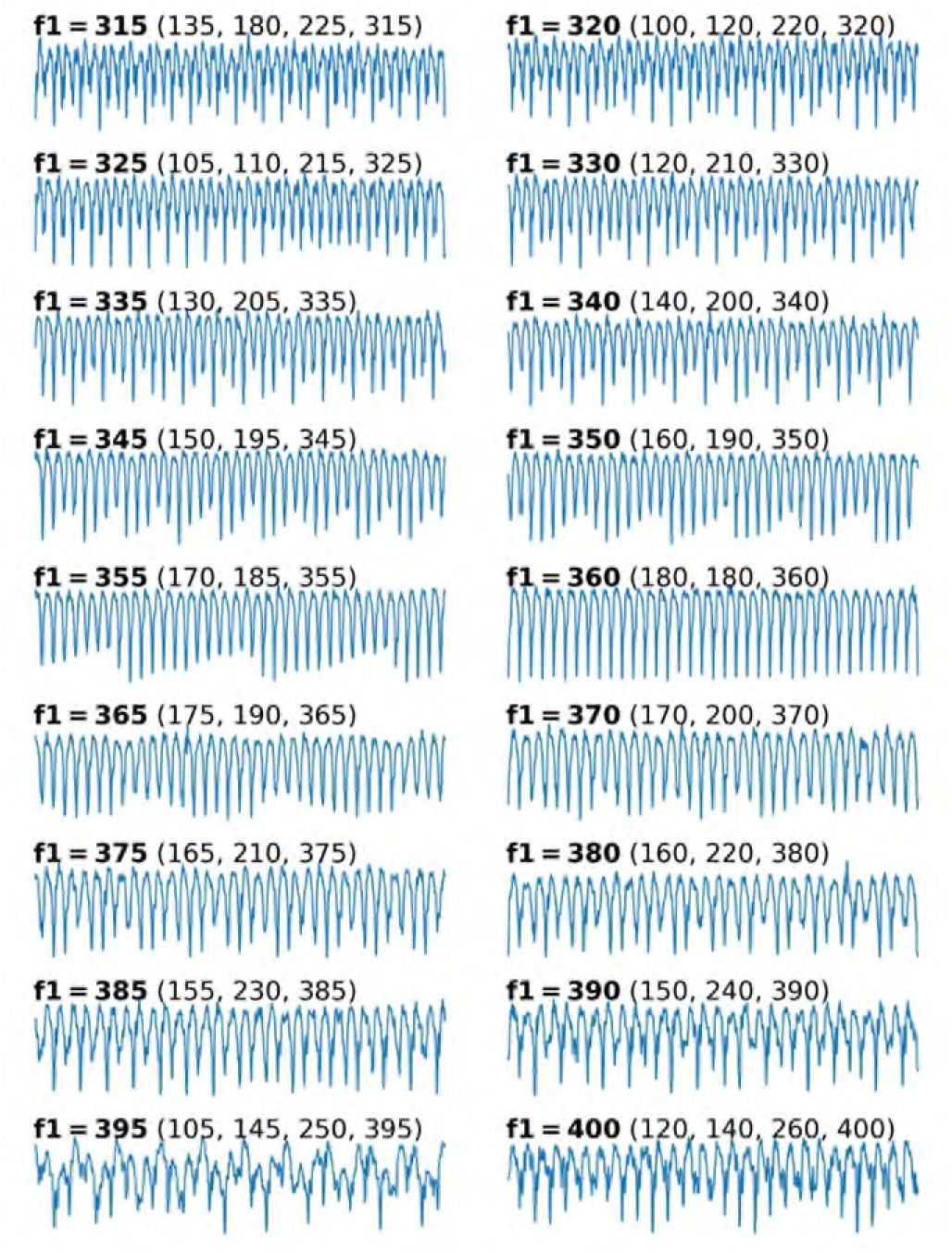
The distortional content of the nerve significantly influences the shape of the recorded local field potential waveform. Shown are examples of nerve responses (blue) to two-tone stimuli with ***f***_2_ fixed at 540 Hz, and ***f***_1_ varying from 315 to 400 Hz (left to right and top to bottom; ***f***_1_ denoted in bold fonts). Distortion products with frequencies between 100 and 400 Hz – the sensitivity range of the mosquito auditory nerve – are displayed in brackets above their corresponding nerve response traces. These data illustrate that the distortional content substantially shapes the overall waveform, as local field potentials reflect the summed activity of multiple neural units, with each distortion frequency recruiting a distinct set of units. Given such waveforms, it is challenging to identify distinct streams of compound action potentials and extract their precisions. An exception is observed for the stimulus pair with ***f***_1_ at 360 Hz and ***f***_2_ at 540 Hz, for which all distortions produced are frequency multiples of 180 Hz and thus all audible components remain phase-locked to each other.

**fig. S7.**
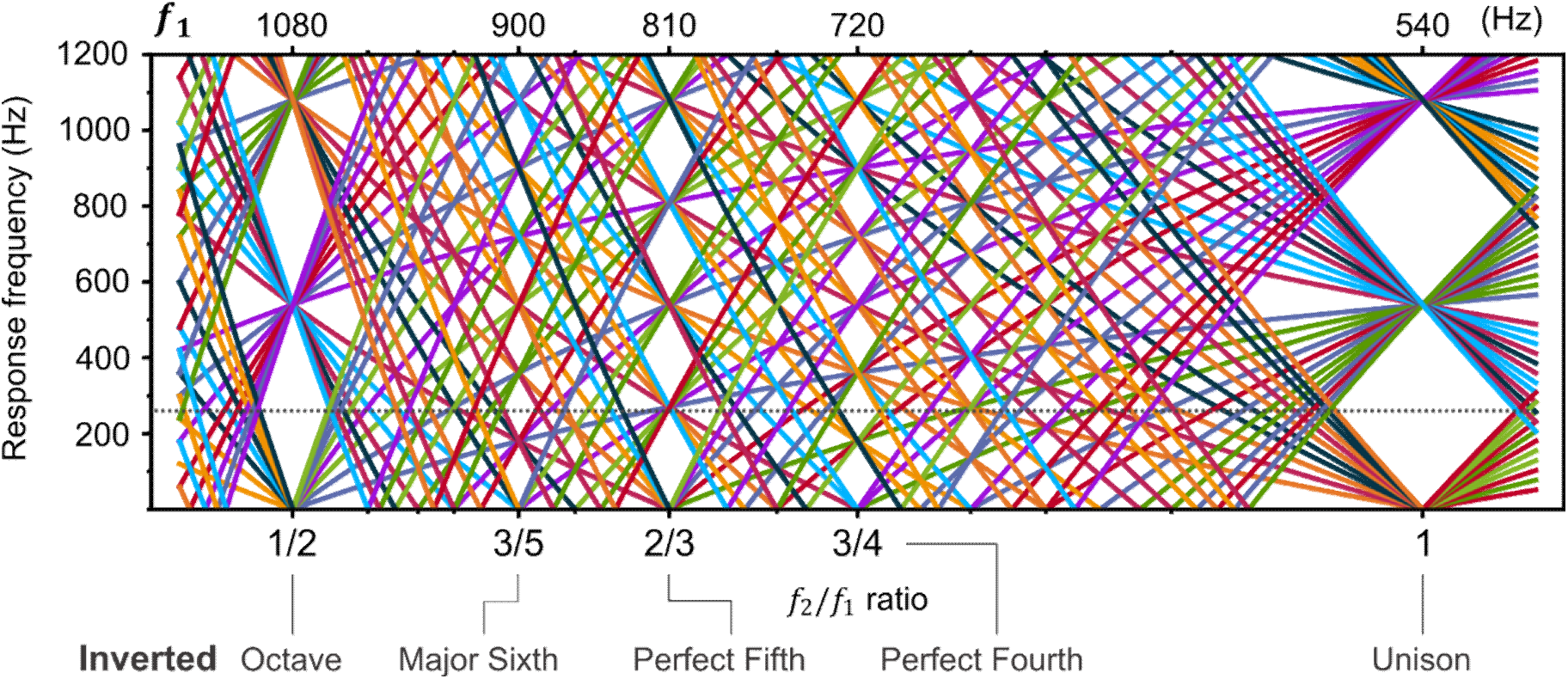
Simulation of two-tone DP families at the inverted musical intervals. While keeping ***f***_2_ fixed at 540 Hz, the ratio is spanned to lower values to show the inverted musical intervals as shown in Fig. 6C. Beyond the inverted octave and unison which are harmonically related to the primary tones, the inverted perfect fifth (2/3) remains the only ratio which enjoys the largest reduction of spectral components. In this case however, DPs are funneled into the ***f***_2_ = 270 Hz subharmonic. The auditory system of quiescent mosquitoes does indeed exhibit heightened nerve responses at 720 ***f***_1_ − ***f***_2_) and 900 Hz (2***f***_2_ − ***f***_1_) as well as at 810 Hz where the two DP families converge (Fig.2B).

**fig. S8.**
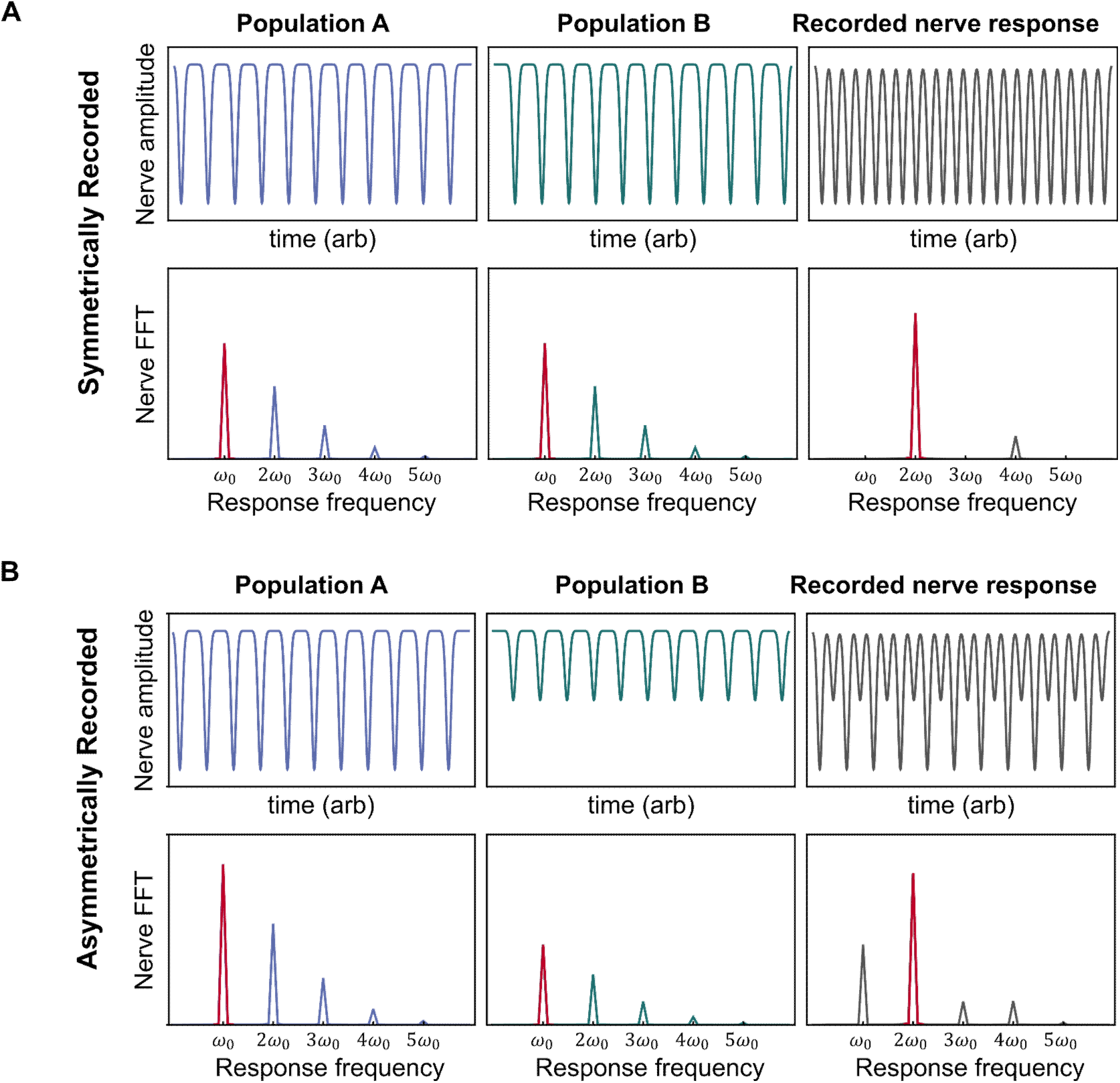
The presence of two identical but physically opposing neuron populations lead to a pronounced 2nd harmonic in the response spectrum. Two neuron populations, A (blue) and B (green), respond to stimulus by firing at a frequency 𝝎_0_ . The two populations are in anti-phase as the flagellar oscillation excites only one population per half-cycle. The spectral composition of their simplified CAP responses is characteristically given by 𝝎_0_ and its harmonics, with the fundamental dominating the spectrum (red). **(A)** If the recording probe is equidistant from the two populations, their relative magnitudes manifest as equal contributors to the nerve response (gray) and the FFT response is effectively described by the second harmonic (2𝝎_0_). **(B)** If the recording probe is closer to population A, the nerve recording yields the summed response from two unequal populations, which adds complexity to the spectrum and no longer follows the characteristic spectra of a spike train. In both cases, however, the 2nd harmonic is dominant regardless of the probe position. The results here are in good agreement with the observed one-tone responses shown in Fig.2A.

**fig. S9.**
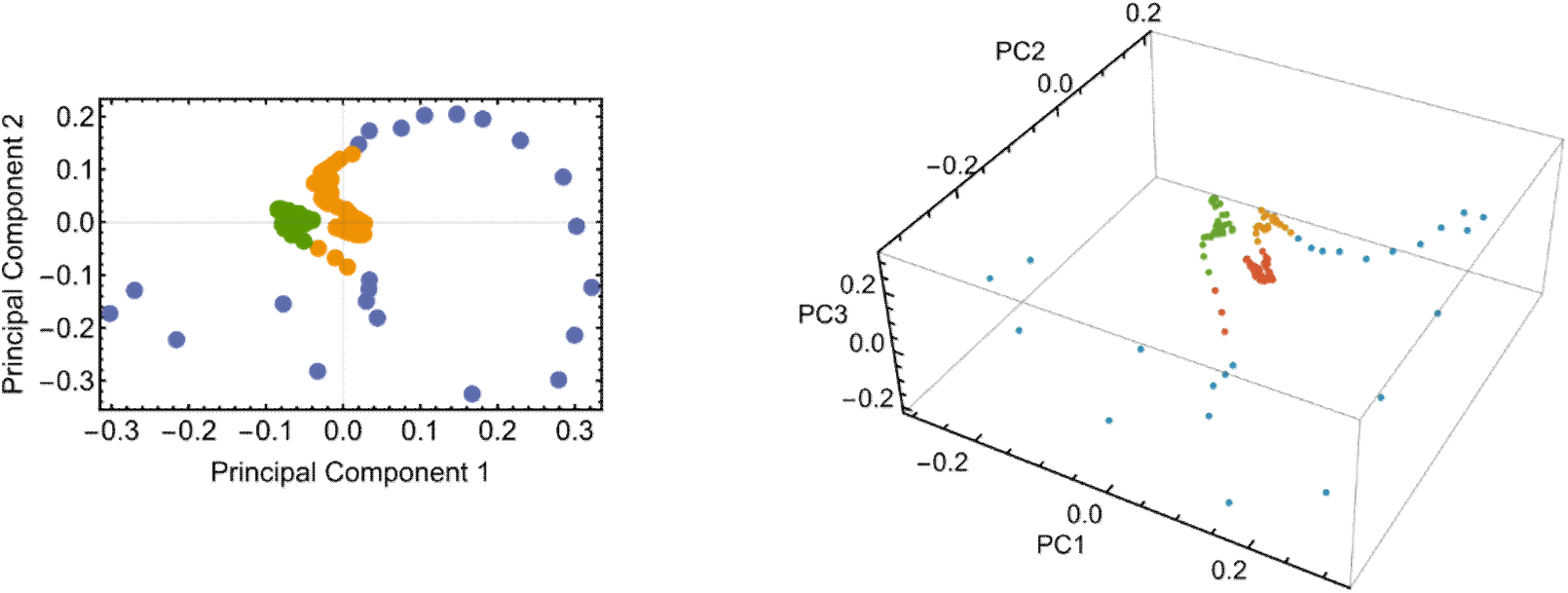
Clustering of animals based on their flagellar mechanical state. The mosquitoes in this study were grouped based on their flagellar mechanical state into three groups: quiescent, weak SSO and SSO. The clustering was done based on principal component analysis applied onto the flagellar envelopes as shown in Fig.1c of the main text. Despite the simplicity of the flagellar envelope, the presence of the SSO – which can vary both in frequency and amplitude – is a strong modulator and dramatically affects the classification into distinct eigenstates. Casting the eigenvectors into two (left), or three (right) dimensions, the animals can be categorized into either three or four classes. A continuous band (in blue) indicates that animals can smoothly cross from each cluster meaning that the SSO amplitude is likely a major factor.

**table S1.** List of DP families in the flagellum or nerve recordings.

|  |  |  |  |  |  |  |  |
| --- | --- | --- | --- | --- | --- | --- | --- |
| A1 | $f_2 - f_1$ | B1 | $2f_1 - f_2$ | C1 | $2f_2 - 3f_1$ | D1 | $3f_2 - 4f_1$ |
| A2 | $f_1 - f_2$ | B2 | $2(f_2 - f_1)$ | C2 | $5f_2 - 3f_1$ | D2 | $2(2f_1 - f_2)$ |
| A3 | $2f_2 - f_1$ | B3 | $3f_2 - 2f_1$ | C3 | $3f_1 - f_2$ | D3 | $4(f_2 - f_1)$ |
| A4 | $f_1 + f_2$ | B4 | $2f_1$ | C4 | $3(f_2 - f_1)$ | | |
| A5 | $3f_2 - f_1$ | B5 | $2(f_1 - f_2)$ | C5 | $3f_1 - f_2$ | | |
| | | B6 | $2(f_2 - f_1)$ | C6 | $3f_1$ | | |
| | | B7 | $2f_1 + f_2$ | C7 | $3(f_2 - f_1)$ | | |
| | | B8 | $f_2 - 2f_1$ | C8 | $f_2 - 3f_1$ | | |
| | | | | C9 | $3f_1 + f_2$ | | |

The typical distortion product families observed under one- and two-tone stimuli.

**table S2.**
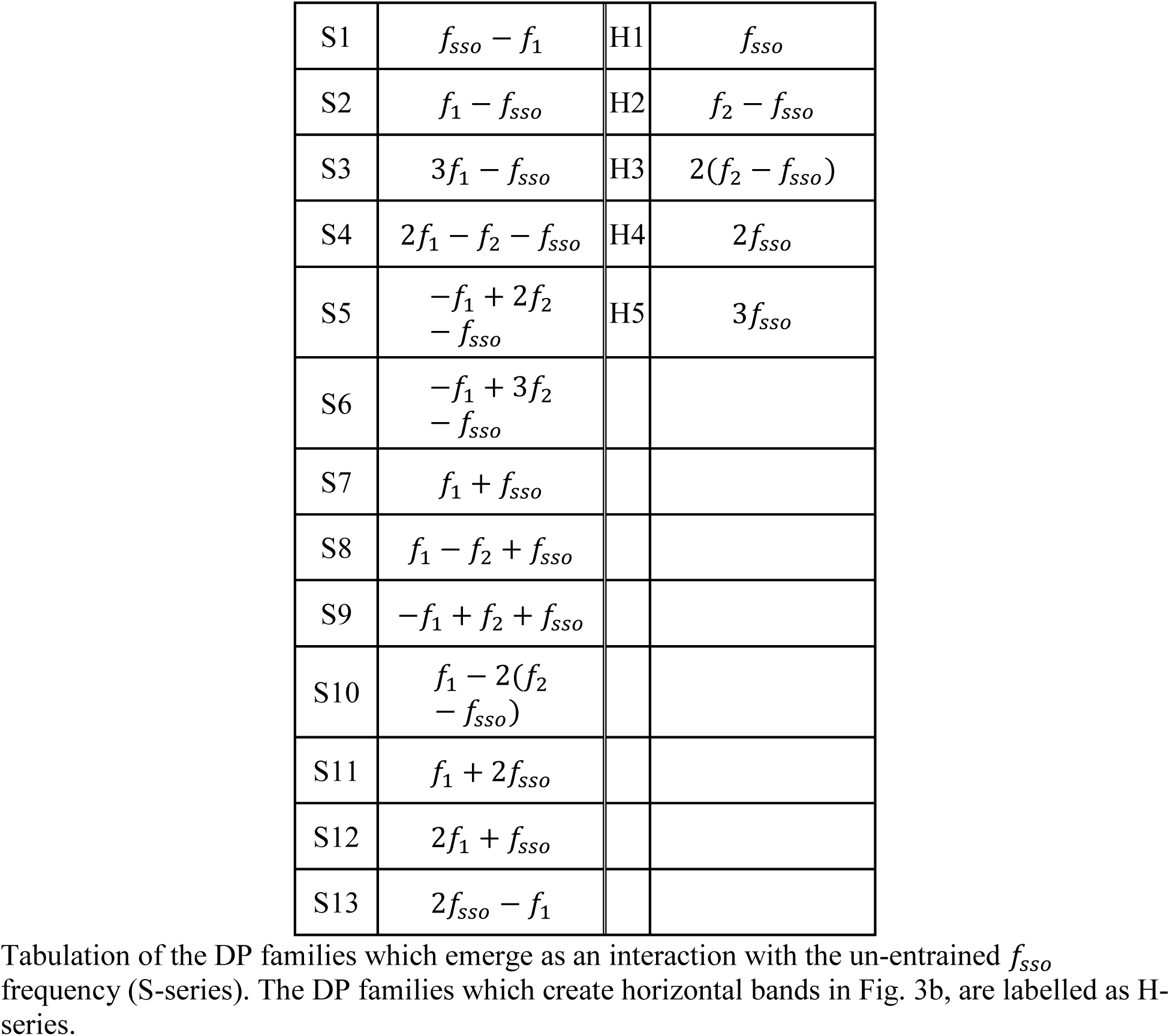
List of DP families containing *f*_sso_.

